# Spatial organization of chromosomes leads to heterogeneous chromatin motion and drives the liquid- or gel-like dynamical behavior of chromatin

**DOI:** 10.1101/2021.05.10.443375

**Authors:** Hossein Salari, Marco Di Stefano, Daniel Jost

**Author notes:** Institute of Human Genetics, Univ of Montpellier, CNRS, Laboratoire de Chromatine et Biologie Cellulaire, Montpellier, France. Corresponding authors (H.S.) and (D.J.).

## Abstract

Chromosome organization and dynamics are involved in regulating many fundamental processes such as gene transcription and DNA repair. Experiments unveiled that chromatin motion is highly heterogeneous inside cell nuclei, ranging from a liquid-like, mobile state to a gel-like, rigid regime. Using polymer modeling, we investigate how these different physical states and dynamical heterogeneities may emerge from the same structural mechanisms. We found that the formation of topologically-associating domains (TADs) is a key driver of chromatin motion heterogeneity. In particular, we demonstrated that the local degree of compaction of the TAD regulates the transition from a weakly compact, fluid state of chromatin to a more compact, gel state exhibiting anomalous diffusion and coherent motion. Our work provides a comprehensive study of chromosome dynamics and a unified view of chromatin motion enabling to interpret the wide variety of dynamical behaviors observed experimentally across different biological conditions, suggesting that the ‘liquid’ or ‘solid’ behaviour of chromatin are in fact two sides of the same coin.

## Introduction

The structural and dynamical properties of the eukaryotic genome inside the cell nucleus play crucial roles in many cell functions, such as gene regulation (van Steensel and Furlong 2019). Over the last decade, high-throughput chromosome conformation capture (Hi-C) experiments have provided valuable information about how genomes organize by measuring the contact frequencies between all pairs of chromatin loci (Lieberman-Aiden et al. 2009). Analyses of Hi-C contact maps in various species and cell types revealed that interphase chromosomes are partitioned at different scales (Rowley and Corces 2018): from topologically-associating domains (TADs) (Dixon et al. 2012) at few hundreds of kilo-basepairs (kbp) to the euchromatic ‘A’ and heterochromatic ‘B’ compartments at Mbp scales and to chromosome territories at the nuclear scale. Complementary to Hi-C, advances in microscopy on fixed cells confirmed the existence of these architectural motifs at the single-cell level (Wang et al. 2016; Boettiger et al. 2016; Szabo et al. 2018; Bolzer et al. 2005) and showed that at the sub-TAD scale, chromatin organizes into clutches of nucleosomes (Ou et al. 2017; Ricci et al. 2015) clustered into nanodomains (Szabo et al. 2020).

Beyond the ‘static’ picture of a layered spatial organization that contributes to genome regulation, more and more experiments on living cells highlighted the ‘dynamic’ nature of chromatin folding and its importance on key biological functions (Tortora et al. 2020; Bystricky 2015; Shaban et al. 2020a). Chromatin mobility has been proposed to impact the dynamics of promoter-enhancer interactions and thus regulates transcriptional bursting (Bartman et al. 2016; Chen et al. 2018), facilitates homology search after DNA damage (Hauer et al. 2017) or participates in the long-range spreading of epigenomic marks (Jost and Vaillant 2018). Chromatin motion is standardly investigated by monitoring the mean-squared displacement (MSD) after a time-lag Δ*t*, that measures the typical space explored by a locus during Δ*t*. Many live-tracking experiments (Nozaki et al. 2017; Ashwin et al. 2019; Germier et al. 2017; Khanna et al. 2019; Shaban et al. 2020b; Zidovska et al. 2013; Barth et al. 2020; Shaban et al. 2018; Bronshtein et al. 2015; Gu et al. 2018; Nagashima et al. 2019; Hajjoul et al. 2013; Socol et al. 2019; Cabal et al. 2006) have shown that the MSD of an individual locus can be interpreted by the same power-law diffusive model *MSD* = *D*Δ*t*^*α*^, where *D* is the diffusion constant and *α* is the diffusion exponent. A wide variety of diffusion constants and exponents have been observed experimentally (**Fig. 1A**) depending on the cell type (Bronshtein et al. 2015), the transcriptional or physiological state of the cell (Germier et al. 2017; Nagashima et al. 2019; Gu et al. 2018) or the presence of DNA damage (Hauer et al. 2017; Amitai et al. 2017; Herbert et al. 2017; Eaton and Zidovska 2020). Strikingly in many situations, the MSD may exhibit different diffusion regimes (i.e. different *α* values) at different time-lag scales. In addition to such heterogeneity across conditions, chromatin motion is also highly heterogeneous inside individual nuclei (**Fig. 1B**,**C**) as observed by genome-wide experiments of chromatin dynamics (Shaban et al. 2020b; Ashwin et al. 2019; Bronshtein et al. 2015) which detected, at a same time, populations of loci with high or low mobility (Lerner et al. 2020; Shaban et al. 2020b; Ashwin et al. 2019). These studies also revealed the presence of spatial chromatin domains of correlated motions (Zidovska et al. 2013; Shaban and Seeber 2020). However, the determinants driving the mobility of individual loci or the formation of such domains are still unclear as some studies associate them with the hetero/euchromatin compartmentalization (Nozaki et al. 2017; Lerner et al. 2020) or cohesin-mediated TAD organization (Ashwin et al. 2019) while others did not observe significant correlation between such heterogeneity and chromatin compaction (Shaban et al. 2020b).

**Fig 1:**
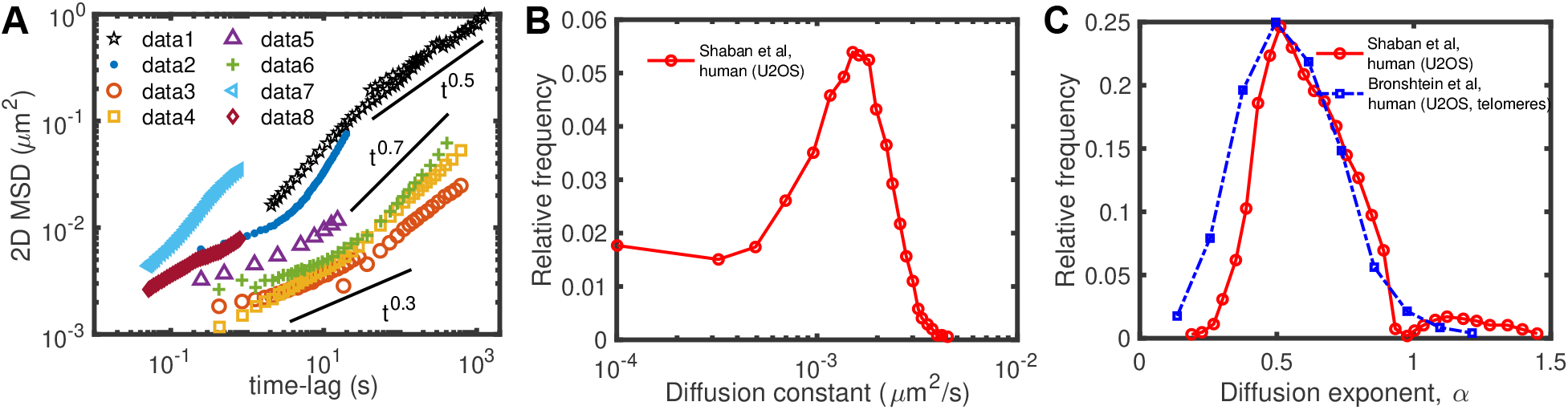
Heterogeneity of chromatin motion. **(A)** Examples of ensemble-averaged mean-squared displacement (MSD) profiles measured experimentally at individual loci for different organisms and cell lines. The different datasets are from: (data1) Mouse pro-B (Khanna et al. 2019), (data2) human MCF-7 (not transcribed gene) (Germier et al. 2017), (data3) human U2OS (centromeres) (Bronshtein et al. 2015), (data4) human U2OS (telomeres) (Bronshtein et al. 2015), (data5) human HeLa cells (Zidovska et al. 2013), (data6) mouse MF (Bronshtein et al. 2015), (data7 and data8) human HeLaS3 (fast loci and slow loci, respectively) (Ashwin et al. 2019). **(B**,**C)** Distributions of diffusion constants (B) and diffusion exponents (C) inferred from the time-averaged MSDs of different loci measured in human U2OS cells (data extracted from (Shaban et al. 2020b; Bronshtein et al. 2015)).

Biophysical modeling has been instrumental in interpreting and predicting the outcomes of live imaging experiments on chromatin motion (Tortora et al. 2020; Di Stefano et al. 2021). Indeed, in classical kinetic theory, the value of the diffusion exponent *α* may be a good indicator of the main underlying physical processes driving the motion of the object under study. For small particles, while standard diffusion is characterized by *α* = 1, subdiffusion (*α <* 1) and superdiffusion (*α* > 1) may indicate constrained or facilitated movement, respectively. For polymers which are large molecules with many internal degrees of freedom, the fixed connectivity along the chain constrains the motion of individual monomers. The Rouse model, a standard polymer theory assuming that mobility is only driven by thermal fluctuations (Doi et al. 1988), thus predicts that the loci of the polymer chromatin should experience a subdiffusive motion with *α* ∼ 0.5, which is the average typical exponent measured experimentally (**Fig. 1**). One can then define a sub-Rousean (*α <* 0.5) and a super-Rousean (*α* > 0.5) diffusion regimes for polymers that may translate additional constraints or forces acting on the monomer motion. Therefore, several decorated Rouse-like models have been developed along the years to suggest that the observed sub-Rousean dynamics may be associated with condensation of chromatin (Shi et al. 2018; Di Pierro et al. 2018) and the super-Rousean regimes with active processes (Chaki and Chakrabarti 2019; Foglino et al. 2019) (see (Tortora et al. 2020) for a review). Dynamical simulations of copolymer models capturing quantitatively the different layers of chromosome organization (Di Pierro et al. 2018; Liu et al. 2018; Shi et al. 2018; Ghosh and Jost 2018; Shukron and Holcman 2017; Shukron et al. 2019) are consistent with an average sub-Rousean regime, with different mobilities between eu- and heterochromatic regions and with correlated motion associated with compartmentalization. In particular, Shi et al associated the experimentally-observed heterogeneity in chromatin motion to the intrinsic glassy dynamics of chromosomes (Shi et al. 2018), while Shukron et al suggested that it emerges from cell-to-cell variability in cross-linking sites (Shukron and Holcman 2017; Shukron et al. 2019).

All these experimental and theoretical works draw a composite - and relatively controversial - picture of how chromatin moves inside cell nuclei during interphase and of how this heterogeneity in motion emerges from fundamental processes and from chromatin architecture. In particular, this has led to two main descriptions of chromatin motion, based on an analogy with materials science (Strickfaden 2021): chromatin behaves like a ‘liquid’ or a ‘fluid’ (Maeshima et al. 2016; Ashwin et al. 2020) pointing to a dynamic and mobile view of chromatin motion; or behaves like a ‘gel’ or a ‘solid’ (Khanna et al. 2019; Strickfaden et al. 2020; Eshghi et al. 2021) highlighting a more constrained dynamics and rigid state.

In order to shed light into this controversy, we investigated how the heterogeneous and anomalous behaviors of chromatin mobility may emerge from first principles using polymer modeling. In particular, we addressed the interplay between the three-dimensional chromosome organization and the different diffusion regimes of chromatin observed experimentally by investigating the dynamics of heteropolymer models that quantitatively describe the chromosome architecture.

## Results

### Quantitative data-driven modeling of 3D chromosome organization

To investigate chromatin motion in situations compatible with experiments, we first developed a data-driven polymer model to quantitatively describe the 3D chromatin organization. We modeled chromatin as a coarse-grained heteropolymer (Fig. 2A). Each monomer, containing 2 kbp of DNA and being of size 50 nm, is characterized by three structural features inferred from Hi-C maps (see **Methods**): its TAD, its compartment (A or B), and, optionally, its anchoring role in CTCF-mediated loops as often observed at TAD boundaries in mammals (Rao et al. 2014; Dowen et al. 2014). The spatio-temporal dynamics of the system is governed by generic properties of a homopolymer (excluded volume and bending rigidity) (Ghosh and Jost 2018) decorated by three types of short-ranged attractive interactions accounting for the heterogeneity of monomer states (see **Fig.2A** and **Methods**): intra- and inter-compartment (*E*_*AA*_, *E*_*BB*_, *E*_*AB*_), intra-TAD with a strength that depends on the local compartmentalization (*E*_*TAD,A*_, *E*_*TAD,B*_), and looping between CTCF anchors (*E*_*loop*_). Note that our approach does not aim to investigate how loops, TADs or compartments emerge from first-principle mechanisms but rather to fold an effective polymer model that captures the main organizational features of chromosomes from Hi-C and serves to study their consequences on chromatin dynamics. Therefore, our model may account for actual attractive forces mediated by chromatin-binding proteins like PRC1 or HP1 (Strom et al. 2017; Larson et al. 2017; Isono et al. 2013; Plys et al. 2019) but also effectively for other mechanisms acting on genome folding like cohesin-mediated loop extrusion (Fudenberg et al. 2016; Sanborn et al. 2015).

**Fig 2:**
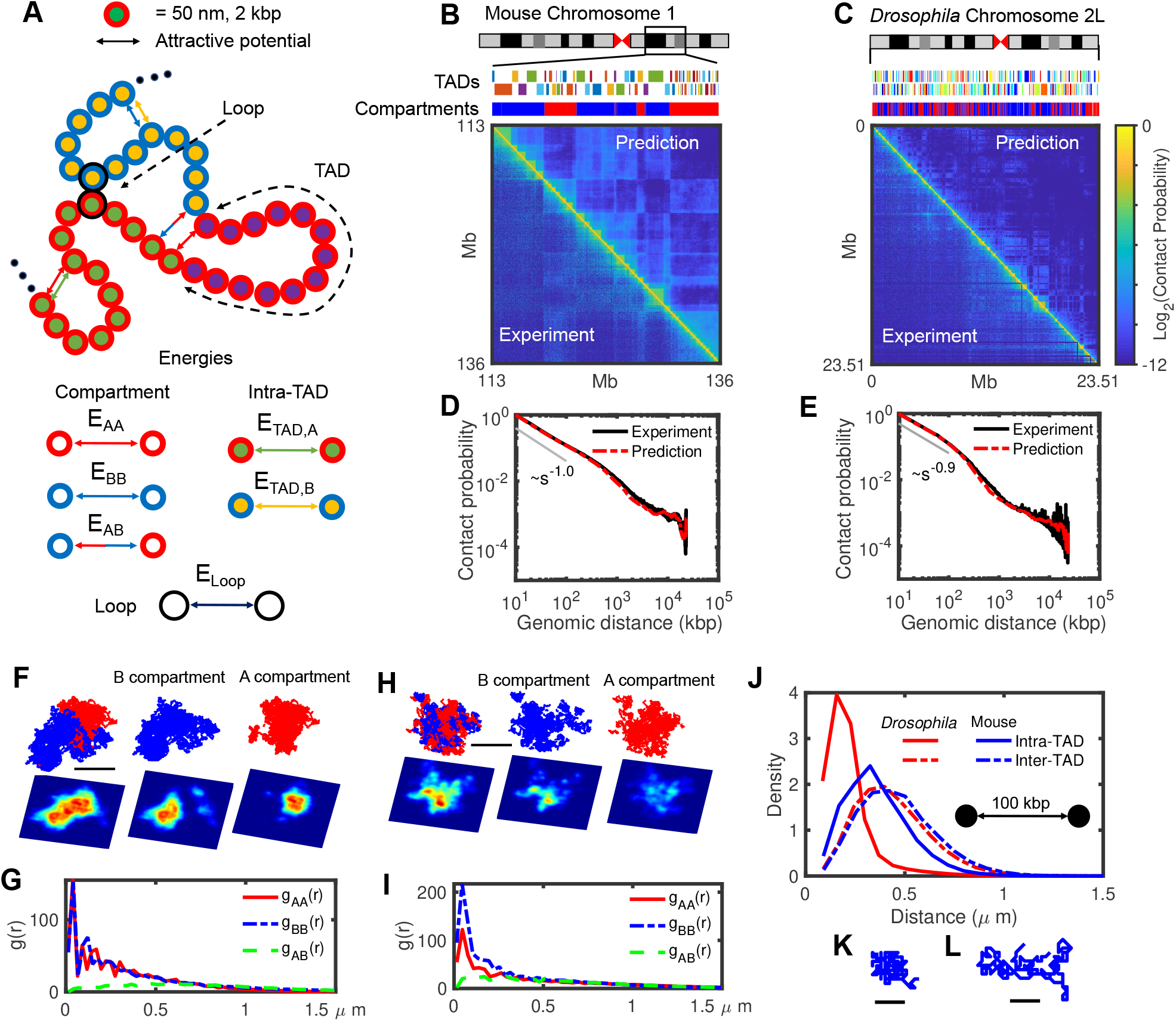
The heteropolymer model and comparison with Hi-C maps. **(A)** Schematic representation of the heteropolymer model with different structural components and their associated interactions. Each monomer hosts 2 kbp of DNA and its diameter is 50 nm. The surrounding red (blue) color indicates A (B) compartment, inner different colors correspond to different TADs, and loop anchors are shown by black outer circles. All six possible attractive interactions are shown by arrows in this cartoon. **(B**,**C)** Visual comparison of predicted Hi-C maps and the experimental ones for 23 Mbp of mouse Chromosome 1 (113-136 Mbp) and *Drosophila* Chromosome 2L. The upper tracks show the corresponding TADs and compartments for each chromosome. **(D**,**E)** Contact probabilities extracted from predicted (red) and experimental (black) Hi-C maps shown in panels (B) and (C), respectively. **(F**,**H)** Typical snapshots for mouse Chr 1 (F) and *Drosophila* Chr 2L (H) in mixed and separate A (red) and B (blue) compartments, with corresponding 2D density plots (the lower panels) with axial resolution of 1,200 nm, lateral resolution of 100 nm and pixel size of 50 nm. Bars represent 1μm of real space. **(G**,**I)** Radial distribution functions (*g*) between A-A (red), B-B (blue) and A-B (green) monomers for mouse (G) and *Drosophila* (I). **(J)** Comparison between intra-(full curves) and inter-TAD (dashed curves) distance distributions of pair monomers separated by 100 kbp along the genome for *Drosophila* (red) and mouse (blue). **(K**,**L)** Typical snapshots of ∼300 kbp long TADs for *Drosophila* (K) and mouse (L). Bars represent 0.25μm of real space.

Next, we optimized the parameters (**Methods**) of the model for two regions of interest (**Table 1**): a 23 Mbp-long portion of Chromosome 1 (113-136 Mbp) in mouse embryonic stem cells (mESC) (**Fig.2B**) and the Chromosome arm 2L in *Drosophila melanogaster* Kc167 female embryonic cells (**Fig.2C**). Our choice for these two cell types aimed to investigate two distinct situations with different TAD and compartment sizes (see the upper tracks in **Fig.2B,C**) and compaction levels but under the same modeling framework. Briefly, we used Hi-C maps as an input to detect TADs, compartments and loop anchors, and thus to define the state of each monomer. Then, by varying the energy interactions, we inferred for each species the parameter set that best predicts the experimental Hi-C (see **Methods** and **Table 2**). As expected, TAD and compartment patterns are qualitatively well described in our predictions (**Fig.2B,C**) as we used that information directly extracted from experiments to build the model. However, in addition, the heteropolymer model was also able, in both cases, to quantitatively reproduce the absolute magnitude of the contact frequencies observed in experimental Hi-C data with an overall high accuracy (correlation of 0.95 for mouse data and 0.87 for *Drosophila*) (**Fig.2B-C**). This goodness of fit, not only captures the average generic decay of contact frequency as a function of their genomic distance (**Fig.2D-E**), but also the structural features of Hi-C maps at different scales, including the intra-TAD and intra-compartment compaction levels (see **Suppl. Fig. S1**).

**Table 1:**
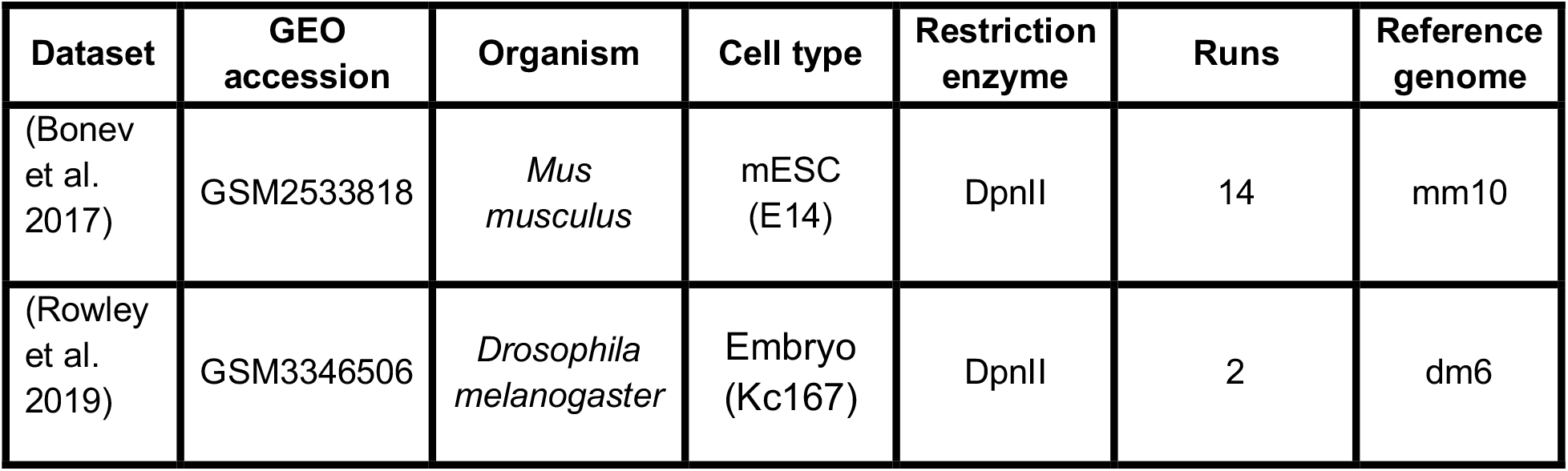
Hi-C datasets used in this study. Data for mouse ES cell and *Drosophila melanogaster* Kc167 were published in (Bonev et al. 2017) and (Rowley et al. 2019), respectively. Dataset, GEO accession number, organism, cell type, restriction enzyme, number of independent runs, and reference genome to map data are given.

**Table 2:**
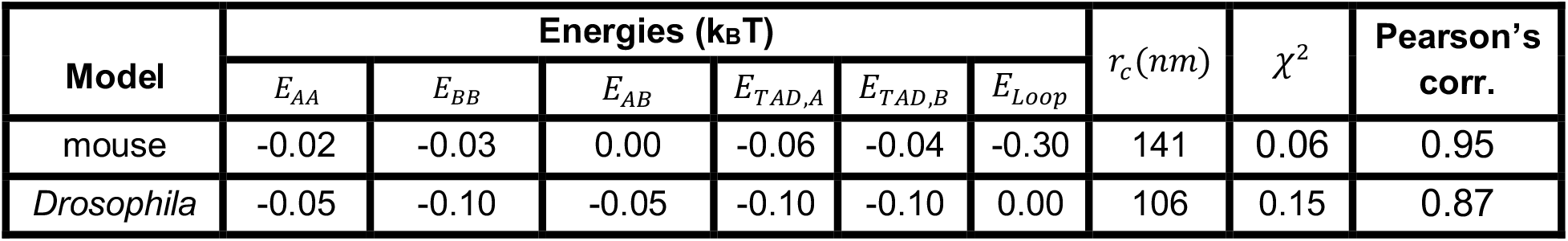
Optimized parameter sets and corresponding Pearson’s correlation and χ^2^ values for mouse Chromosome 1:113-136 Mbp and *Drosophila* Chromosome 2L.

| Model | Energies (k<sub>B</sub>T) | | | | | | $r_c$ (nm) | $\chi^2$ | Pearson’s corr. |
| --- | --- | --- | --- | --- | --- | --- | --- | --- | --- |
| | $E_{AA}$ | $E_{BB}$ | $E_{AB}$ | $E_{TAD,A}$ | $E_{TAD,B}$ | $E_{Loop}$ | | | |
| mouse | -0.02 | -0.03 | 0.00 | -0.06 | -0.04 | -0.30 | 141 | 0.06 | 0.95 |
| *Drosophila* | -0.05 | -0.10 | -0.05 | -0.10 | -0.10 | 0.00 | 106 | 0.15 | 0.87 |

At large scales, the checkerboard-like patterns observed in Hi-C maps suggest that chromatin compartments are spatially segregated. Typical configuration snapshots from our simulations in *Drosophila* **(Fig.2F)** and mouse **(Fig.2G)** indeed illustrate the relative organization of A and B compartments. We quantified this by computing the radial distribution functions (**Fig.2H-I**) *g*_*AA*_(*r*), *g*_*BB*_(*r*) and *g*_*AB*_(*r*) that capture the probabilities to find a monomer of a given compartment at a given distance *r* from a monomer of the same (*g*_*AA*_(*r*),*g*_*BB*_(*r*)) or of a different (*g*_*AB*_(*r*)) compartment. In *Drosophila*, we observed that the B compartment is locally more compact than A (*g*_*BB*_(*r*)>*g*_*AA*_(*r*) for small *r*, **Fig.2H**), while compaction is similar in both compartments in mouse (**Fig.2I**). We also noticed that the A and B compartments are more segregated in mouse, with *g*_*AB*_(*r*) equating *g*_*AA*/*BB*_(*r*) around r*≃700 nm, than in *Drosophila* (r*≃300 nm), resulting in part from the larger genomic size of compartments in the mouse case (see the upper tracks in **Fig.2B-C**).

At the TAD-scale, structural properties are also strongly cell type- and compartment-dependent. On average, TADs in mice (median size ∼120 kbp) are longer than in fly (median size∼40 kbp) (**Fig 2B** and **2C** upper tracks). Globally, TADs in *Drosophila* are more compact than in mice (two typical snapshots of TADs with same size are drawn in **Fig.2K-L**) with a relatively smaller ratio of intra-versus inter-TAD distances of *Drosophila* compared to mice (**Fig.2J**). Similar to the condensation of compartments (**Fig.2H-I**), we also observed that TADs in the A compartment are less compacted than those in B for *Drosophila* (Cattoni et al. 2017; Szabo et al. 2018; Lesage et al. 2019), while being more open and having similar compaction level in mouse (Finn et al. 2019) (**Suppl. Fig. S1G**,**H**).

### Chromatin dynamics is strongly heterogeneous and locus-dependent

Having in hand quantitative polymer models capturing the main structural features of chromosome organization, we investigated the dynamical properties of chromatin predicted by such models. As a reference, null model, we also simulated the dynamics of a simple homopolymer where all compartment-, TADs-, or loop-based interactions were switched off (see **Methods**). One standard observable to probe the local chromatin mobility is the mean-squared displacement (MSD) of individual loci as a function of time. Experimentally, this is typically done either by tracking single fluorescently-labelled loci (Cabal et al. 2006) or by monitoring the dynamics of the nuclear local densities of stained histones (Zidovska et al. 2013), during ∼10-30 seconds. From these experiments, trajectories of individual loci in single cells can be extracted and analyzed to compute two types of MSD: time-averaged and ensemble-averaged MSDs.

From each single trajectory, time-averaged MSDs can be estimated with 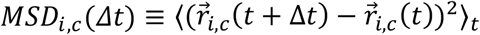 where 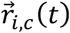 is the position vector of a given locus *i* at time *t* for a trajectory *c* and ⟨⋯ ⟩_*t*_ is the time-averaging along the trajectory *c* (i.e., averaged over *t*). These MSDs can be analyzed by fitting them with a single power-law 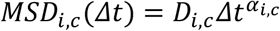 (see **Methods**). Analysis of live experiments have shown high variability in the diffusion constant *D*_*i,c*_ and exponent *α*_*i,c*_ at the single-cell level in many species and for many loci (see examples in **Fig.1B,C**) (Zidovska et al. 2013; Shaban et al. 2020b; Nozaki et al. 2017; Ashwin et al. 2019).

By averaging over all the trajectories *c* for the same locus *i*, ensemble-averaged MSDs can be computed with 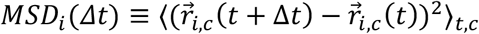. Examples extracted from various single-locus tracking experiments are given in **Fig. 1A** (Hajjoul et al. 2013; Socol et al. 2019; Germier et al. 2017; Bronshtein et al. 2015; Gu et al. 2018). *MSD*_*i*_(Δ*t*) can then be fitted by power-law-like functions 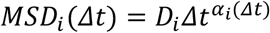 where the diffusion exponent *α*_*i*_ may now depend on the time-lag Δ*t* (see **Methods**). Experiments exhibit a wide variety of sub-diffusive *α*_*i*_-values depending on the locus, species or transcriptional state, with sometimes crossovers between different regimes (see examples in **Fig. 1A**).

To try to understand the origin of this heterogeneity, we first estimated with our simulations the time-averaged MSD for all the monomers in the *Drosophila* and mouse cases over 30 seconds-long trajectories (**Fig.3A** and **Suppl. Fig. S2**,**3**). Distributions of *α*_*i,c*_ are broad (**Fig.3B**) and similar in both cases. The diffusion constants *D*_*i,c*_ are also dispersed (**Fig.3C** and **Suppl. Fig. S3**) with a bimodal distribution for *Drosophila*, exhibiting a population of slowly diffusing trajectories. Our models thus well predict qualitatively the shapes and large variabilities of the distributions of diffusion exponents and constants observed experimentally (**Fig.1B,C**). However, this strong variability is also present in the null - homopolymer - model (**Fig.3A-C**) at the same degree as in the mouse case. This result suggests that part of the heterogeneity observed in time-averaged MSD does not stem from the multi-scale organization of chromosomes in TADs or compartments, but rather from a finite-size effect due to the limited duration of the monitored trajectories to measure the MSD (**Suppl. Fig.S2**). To mitigate this confounding factor and focus on the role of structural heterogeneities on dynamical variabilities, we next computed the ensemble-averaged MSD of all monomers over ∼2000, ∼1 hour-long, trajectories (**Fig.3D-F**). In the homopolymer model (**Fig 3D**), we observed as expected a uniform (Rousean) behavior for all monomers with *α*_*i*_(Δ*t*) ≈ 0.5 at short time scales (Δ*t*<10 s) and *α*_*i*_(Δ*t*) ≈ 0.4 at longer time scales (Δ*t*>100 s), typical of crumpled polymers (Liu et al. 2018; Ghosh and Jost 2018; Tamm et al. 2015). For mouse and *Drosophila* chromosomes, simulations predicted heterogeneous, locus-dependent MSDs (**Fig 3E and 3F**). The distributions of diffusion constants are broad, implying a large spectrum of loci mobilities (**Fig.3G**). This is particularly visible in the *Drosophila* case (**Fig.3G**) where mobility may vary by up to 3-fold between two monomers. In *Drosophila*, we also predicted the distribution to be multimodal with 2 main peaks at low and high mobility, which is fully consistent with experiments (Gu et al. 2018; Shaban et al. 2020b; Lerner et al. 2020). The distributions of diffusion exponents per locus *α*_*i*_(Δ*t*) at different time scales (**Fig.3H-I**) also support the strong heterogeneities observed in diffusion behaviors. While at short time scales (Δ*t*< 3 s) exponents are rather homogeneous (*α*_*i*_ ≈ 0.5), at larger time scales (Δ*t*> 10 s) *α*_*i*_ becomes highly locus-specific and may strongly vary as a function of time. For example, at Δ*t*=1000 s, *α*_*i*_ varies between 0.25 and 0.65 in *Drosophila* and between 0.3 and 0.5 in mice (**Fig.3I**). This broad range of values, including both sub- (*α*_*i*_<0.5) and super- (*α*_*i*_>0.5) Rousean exponents, is consistent with the large discrepancy observed experimentally in *α*_*i*_ across loci and conditions (**Fig. 1A**).

**Fig 3:**
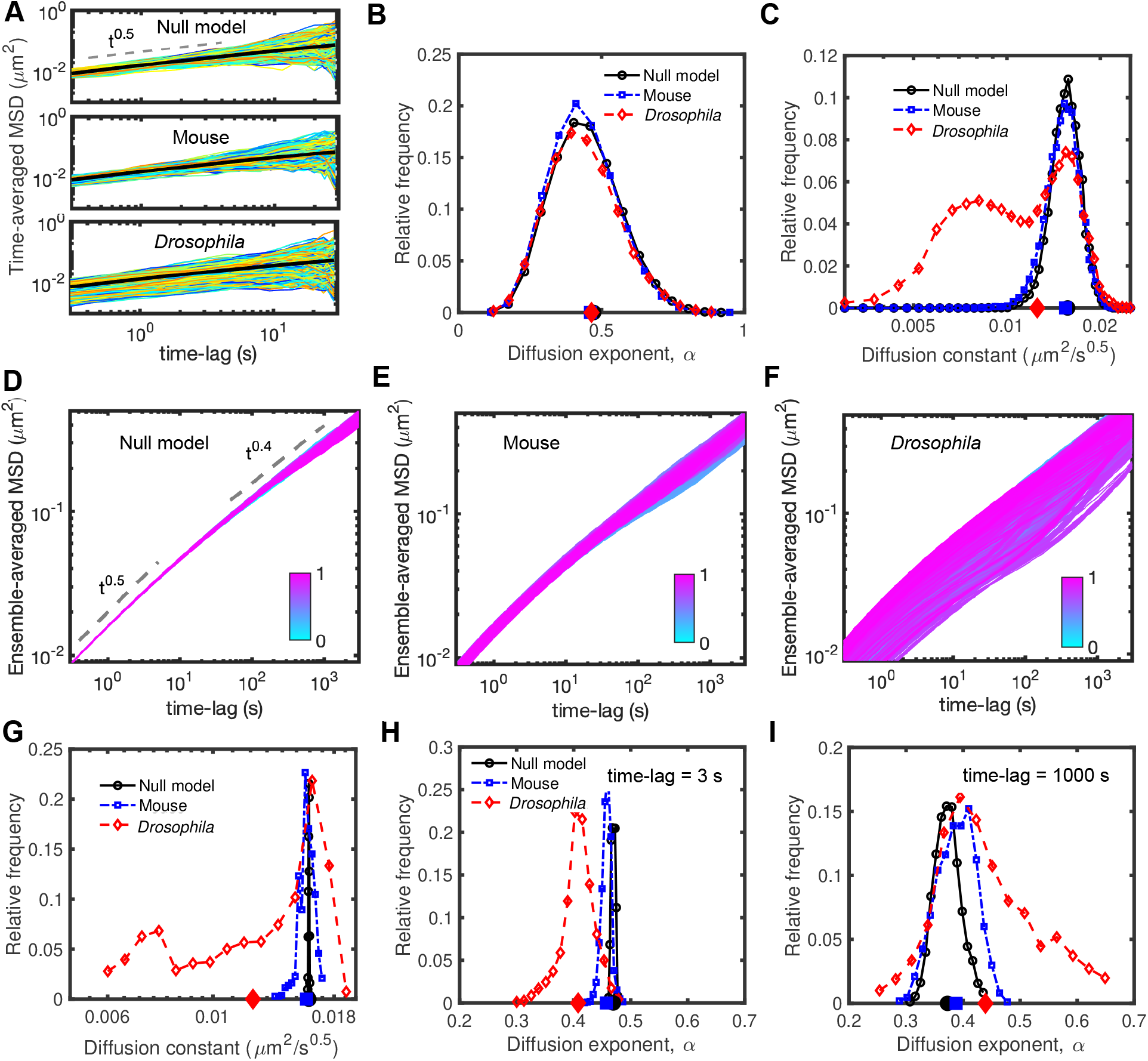
Dynamic properties of simulated chromosomes. **(A)** Time-averaged *MSD*_*i,c*_ of single trajectories of length 30 s sampled every 0.3 s for null (upper panel), mouse (middle panel) and *Drosophila* (bottom panel) models. **(B)** Distribution of the diffusion exponents α_*i,c*_ extracted from the time-averaged MSD curves given in panel (A). **(C)** Distribution of the diffusion constants *D*_*i,c*_ for the time-averaged MSDs in (A) having an exponent α_*i,c*_ ≈ 0.5. Distributions for other α_*i,c*_ values are given in **Suppl. Fig. S3**. In (B,C), average values of the distributions are shown on the horizontal x-axis. **(D**,**E**,**F)** Ensemble-averaged (over all trajectories) *MSD*_*i*_ of all genomic loci for null (D), mouse (E) and *Drosophila* (F) models. To discard trivial positional effects, we exclude the last 50 monomers at the two ends of the polymers. Individual ensemble-averaged MSDs were colored from cyan (first monomer) to magenta (last monomer). **(G**,**H**,**I)** The distributions of the diffusion constant *D*_*i*_ (G) and of the diffusion exponent α_*i*_(Δ*t*) for short (H, Δ*t* = 3 *s*) and large (I, Δ*t* = 1000 *s*) time scales, extracted from ensemble-averaged MSD curves in panels (D-F). The average values of the distributions are shown on the horizontal axes.

**Fig 4:**
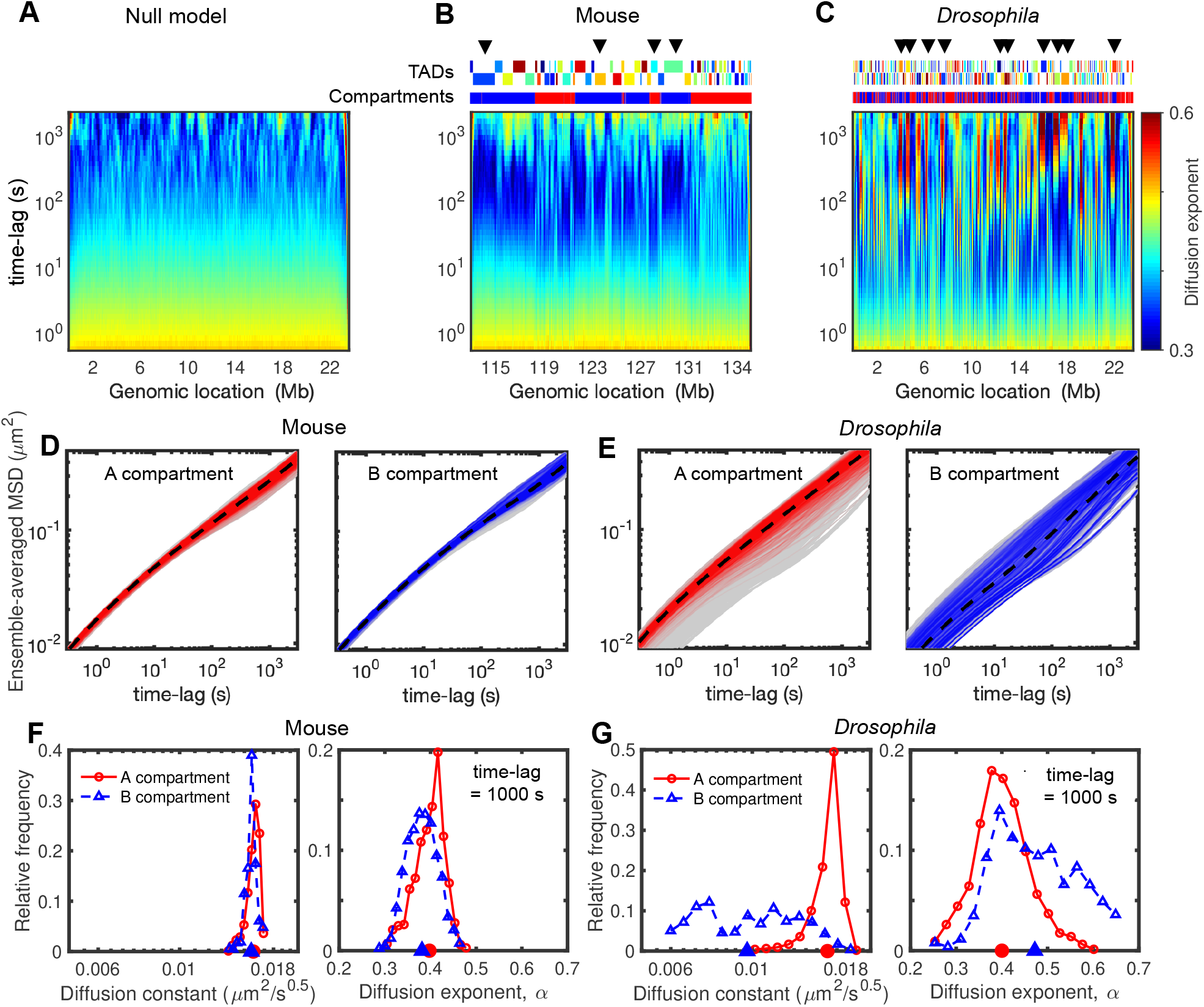
Heterogeneity in dynamics and its relation with compartments. **(A-C)** Time-evolutions of diffusion exponents along the genome for null model (A), mouse (B) and *Drosophila* (C), calculated by the derivatives of logarithmic MSD curves (**Fig. 2G-2I** and **Methods**). The upper tracks in (B) and (C)) show the corresponding TADs and compartments, and arrows highlight some regions with anomalous behaviors (nonmonotonic evolutions). **(D)** MSD curves of the monomers in A (left panel) and B (right panel) compartments in the mouse case. The dashed black curves show the average MSDs over all loci in the same compartment, and gray shaded areas are MSDs of all monomers. **(E)** As in (D) but for the *Drosophila* model. **(F)** The distributions of diffusion constant (left panel) and exponent (right panel) at 1000 s time-lag for the monomers in A (red) and B (blue) compartments in the mouse case. Their average values are indicated on the horizontal axis. **(G)** As in (F) but for *Drosophila* simulations.

### Dissecting the role of compartments and TADs on chromatin motion

The degree of dynamical heterogeneity predicted by the model can only arise from the different interactions driving the TAD and compartment formation. Fig.4 A-C illustrate the evolution of the exponent *α*_*i*_ as a function of the time-lag Δ*t* along the genome. As expected, we observed for the null model an overall homogeneity in dynamical parameters over time (**Fig.4A**). For the mouse and *Drosophila* models, we measured, instead, a strong heterogeneity and locus-dependency along the genome (**Fig.4B,C**). Loci of the same TAD or compartment may have similar time-evolutions of *α*_*i*_ (see also **Suppl. Fig. S4**), suggesting a coupling between dynamics and the different layers of genome folding. Many regions (**Fig.4B,C**) exhibit an anomalous and nonmonotonic behavior: *α*_*i*_ is ∼0.5 at short time scales, decreases to ∼ 0.*A* (sub-Rousean regime) at intermediate time scales, increases to ∼ 0.*B* (super-Rousean regime) and then retrieves a standard crumpled polymer-like behavior (*α*_*i*_ ∼ 0.4) at very large time scales. Such crossovers between sub- and super-Rousean regimes or between super-Rousean and normal regimes have also been observed experimentally (see **Fig.1A**).

### Chromatin mobility is reduced in compact compartments

To quantitatively associate such anomalous behaviors with structural features, we first separately plotted the MSD for loci in active (A) and repressive (B) compartments (**Fig.4D,E**). In mice (**Fig.4D**), both compartments have a similar - weak - degree of heterogeneity with comparable diffusion constants and exponents (**Fig.4F**). This confirms that the dynamical heterogeneity and difference between eu- and heterochromatic loci are usually weak in highly plastic cells as experimentally observed in mESCs (Nozaki et al. 2017) and in human cancerous U2OS cells (Shaban et al. 2020b). In *Drosophila* (**Fig.4E**), loci in the A compartment are on average more mobile than those in the B compartment with a mean increase in mobility of ∼60% for A monomers (**Fig.4G** left panel). In fact, the A monomers correspond to the high mobility peak observed in **Fig.3G**. This is consistent with the observation that differentiated cells (like Kc167) exhibit significant differences in mobility between active and repressive regions (Nozaki et al. 2017; Lerner et al. 2020). We also observed that the distribution of exponents is much broader for B than for A monomers (**Fig.4G** right panel), suggesting globally more heterogeneity and more loci with anomalous behaviors in the B compartment. However, not all the loci in B have anomalous dynamics (**Fig.4E** right panel) and all types of behaviors present in the general population of monomers (**Fig.3F**) are observed in the B compartment. This suggests that compartmentalization per se is not the main driving force of heterogeneity in our model.

### Anomalous behavior is associated with TAD compaction

We thus reasoned that TADs may be a good suspect for driving the anomalous, nonmonotonic diffusion observed in our heteropolymer models. To address this, we first selected two TADs, one from *Drosophila* (size ∼ 480 kbp, 50% reduction in volume compared to a region of similar size in the null model) and one from mouse (size ∼ 1.52 Mbp, 30% volume reduction) with different degrees of compaction and different sizes (**Fig.5A,C**). For the mouse TAD (**Fig.5A,B**) which is bigger but less compact, anomalous behavior is weaker and most of the monomers follow a null model-like behavior. For the *Drosophila* TAD (**Fig.5C**) which is smaller but more compact, we observed that all loci inside the TAD move with anomalous dynamics (**Fig 5D**). Since one main difference between mouse and *Drosophila* heteropolymer models is the intra-TAD strength of interaction (see **Table 2** and **Fig.2J**), these observations point towards an important role played by the intra-TAD compaction level on driving anomalous behaviors. Moreover, by visually comparing the local Hi-C maps and the time-evolutions of the diffusion exponent **Fig 5A**,**C**), we also remarked that the small neighboring TADs (quoted #3 in **Fig.5A,C**) exhibit null-like dynamics (**Fig 5B,D**), while being well formed, suggesting that TAD length might also be an important parameter for anomalousness.

**Fig 5:**
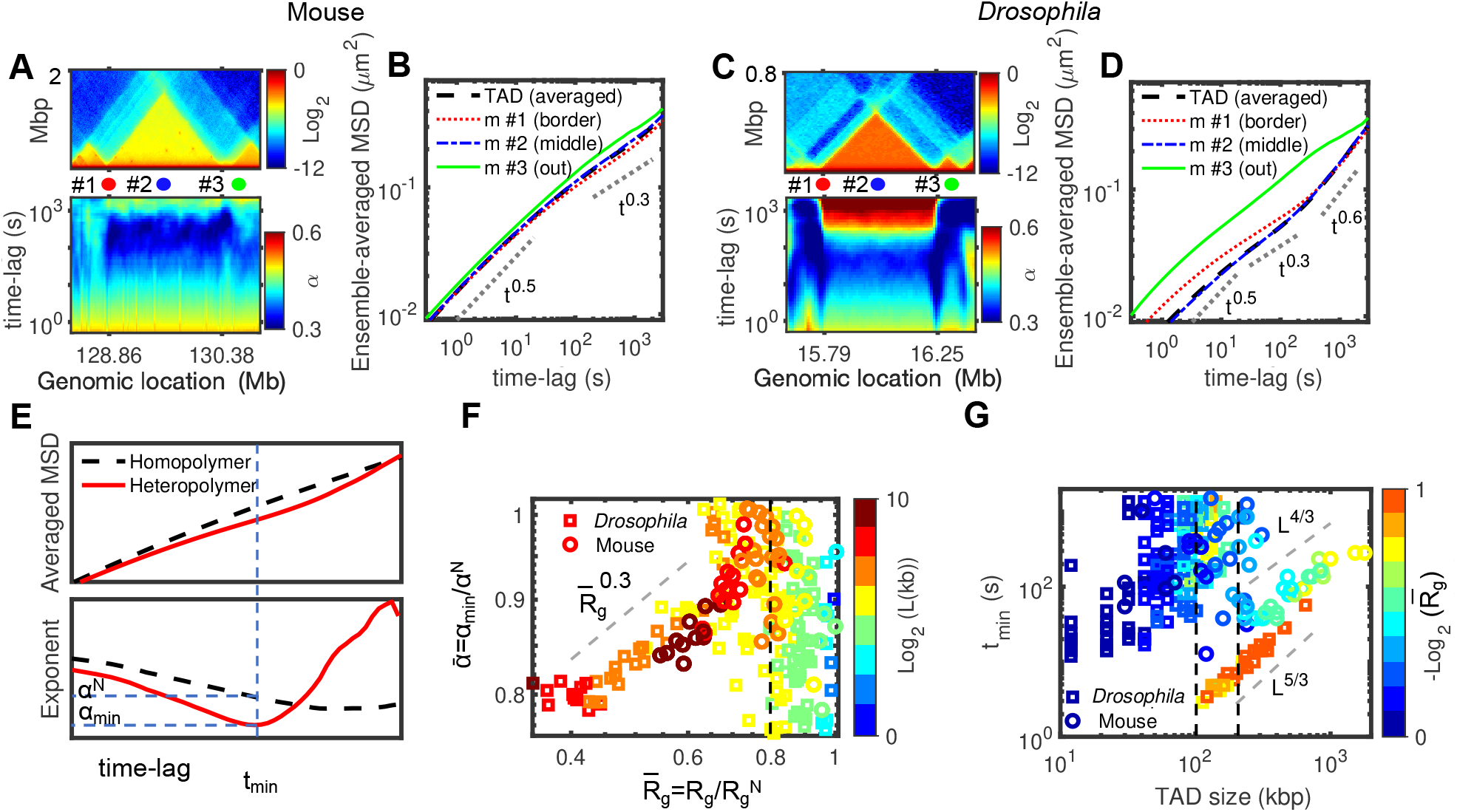
Compaction leads to anomalous behavior. **(A)** Comparison of the predicted Hi-C map (top) and the corresponding time-evolution of the diffusion exponent (bottom) for the mouse Chr 1:128.36-130.88 Mbp region, which includes a 1.52 Mbp-long TAD. **(B)** MSD for three monomers annotated in panel (A) (red: m #1 at the border of TAD, blue: m #2 at the middle of the TAD, green: m #3 outside the TAD) and average MSD for all the monomers inside the TAD. **(C-D)** As in (A-B) but for the *Drosophila* Chr 2L:15.64-16.4 Mbp region, which includes a 460 kb-long TAD. **(E)** Typical average MSDs for a TAD predicted by the homopolymer and heteropolymer models (top), and the time-evolution of the diffusion exponent (bottom). For the heteropolymer curve, we computed the minimum exponent *α*_*min*_, the corresponding time *t*_*min*_, and the expected diffusion exponent *α*^*N*^ in the homopolymer model at *t*_*min*_. **(F)** 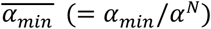 against 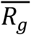 for *Drosophila* and mice. The symbols are colored by the TAD length, *L*. **(G)** *t*_*min*_ as a function of TAD length for *Drosophila* and mice. Symbols are colored by the compaction level of TADs 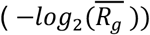.

**Fig 6:**
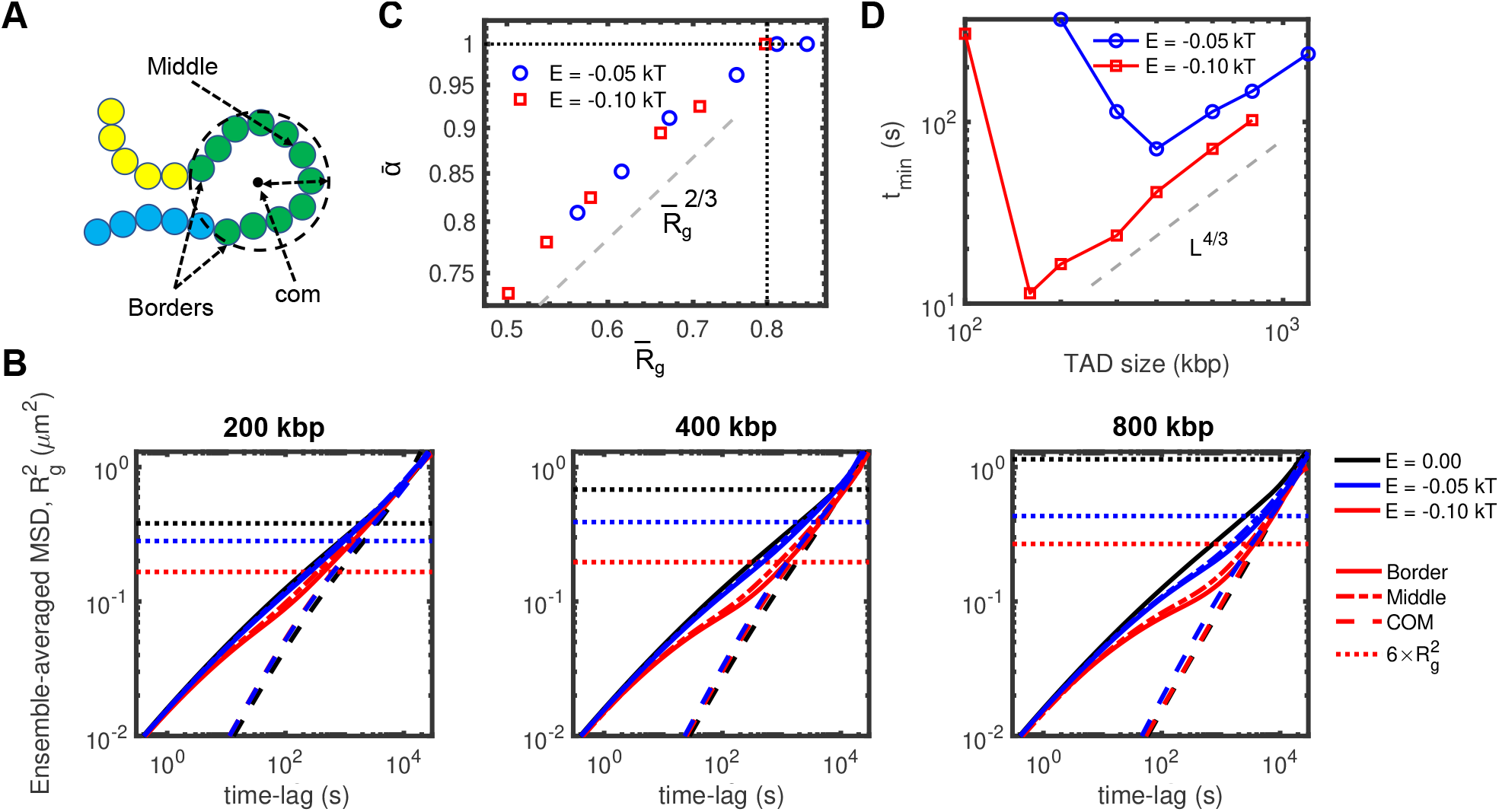
Anomalous behavior in uniform models. **(A)** Cartoon to depict ‘middle’ and ‘border’ monomers, along with the center of mass and the radius of gyration of a TAD. **(B)** Comparison between MSDs of the monomers in the middle (mid) and borders (bor) of TAD and of the center of mass of the TAD (com) (see panel A) from uniform heteropolymer models with TAD length of 200 kbp (left), 400 kbp (middle) and 800 kbp (right) for two different intra-TAD strength of interaction (black: 0kT/null model, blue: -0.05kT, red: -0.1 kT). The spatial sizes 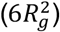 for the different strengths of interaction are shown as dotted lines. **(C**,**D)** 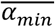 as a function of 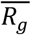 and *t*_*min*_ as a function of the TAD length *L* for the uniform models and for different strengths of interaction.

To quantify this, we considered, for each TAD, 3 ‘structural’ quantities: *L*, the TAD length; *R*_*g*_, its radius of gyration that captures its typical 3D size; 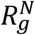, the radius of gyration for a domain of length *L* in the null model. Strongly compacted TADs are characterized by low values for 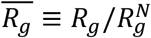. And we introduced 3 ‘dynamical’ observables (**Fig.5E**): *α*_*min*_, the minimum diffusion exponent value over time computed from the average MSD of monomers inside the same TAD; *t*_*min*_, the time when *α*_*min*_ is reached; *α*^*N*^ the exponent of the null model at *t*_*min*_. The strength of the anomalous behavior is thus captured by the ratio 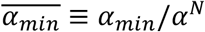 (low ratios *<* 1 corresponding to strong anomalousness), and its duration by *t*_*min*_.

We found that 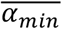 is an increasing function of 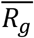 for 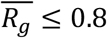 while we do not observe a clear-cut dependence of 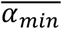 on *L* (**Fig.5F**). *Drosophila* and mouse TADs follow the same scaling law 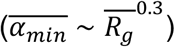, suggesting that compaction level is the main driver of the strength of the anomalous behavior. For weakly compacted domains 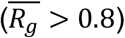 or small TADs, the value of 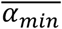 is less well defined. We also observed that, beyond a critical size (∼100 kbp for *Drosophila* and ∼200 kbp for mouse), *t*_*min*_ evolves as ∼ *L*^5/3^ for the *Drosophila* case and ∼ *L*^4/3^ for the mouse case, the role of 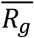 being less clear (**Fig.5G**). These results show that the duration of the anomalous, nonmonotonic diffusion in heteropolymer models depends on the TAD length and compaction, longer domains being impacted for longer times. As observed for 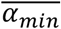, we found that *t*_*min*_ values for small TADs are very dispersed without significant correlation with the TAD length.

### Anomalous behavior emerges from a crossover towards collective motion

The existence of anomalous behaviors with MSDs exhibiting transitions between different - *a priori* opposite (sub- vs super-Rousean) regimes is thus strongly associated with TAD compaction. To go deeper in the analysis of this association and to better understand how these transitions emerge from compaction, we introduced toy heteropolymer models (see **Methods**) where we can independently play with the TAD length and intra-TAD strength of interaction. We considered simplified ‘uniform’ models where genomes are partitioned into adjacent TADs of the same size (see **Methods** and **Suppl. Fig. S5**). For given TAD length and intra-TAD strength of interaction, all monomers of the chain have very similar MSDs (**Fig.6A,B**), with only a weak positional effect translating the bead positioning inside the TAD, consistent with our observations made on more ‘complex’ models (**Fig.5B,D**). As expected, emergence of anomalous behavior occurs for larger TADs and stronger interactions (**Fig.6B** and **Suppl. Fig S6**) where compaction starts to be substantial (**Suppl. Fig. S7**). Merging all the investigated TAD lengths and strengths of interaction together, we confirmed that *t*_*min*_ increases with *L* (**Fig.6D**) and that below a given critical compaction level 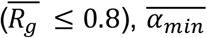 is an increasing function of 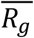 (**Fig.6C**). The dependency is steeper 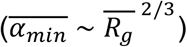 in the uniform models than for the more heterogeneous mouse and *Drosophila* cases. In the uniform models, for small and weakly compacted TADs, we observe 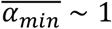. This suggests that the variability of 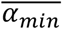 values observed at this compaction regime 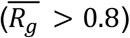 in the heterogeneous *Drosophila* or mouse models (**Fig.5F**) reflects in fact the influence of neighbor - more compacted - TADs on these - normally weakly-impacted - domains (**Suppl. Fig. S8A-C, S9**). By introducing A/B compartmentalization in these uniform models (**Suppl. Fig. S8D-F, Suppl. Fig S10**), we confirmed that more compact compartments show indeed reduced mobility but that the anomalous behavior mostly emerges from the formation of local interaction domains like TADs.

By plotting the MSD of the center of mass of TADs in the uniform models (*MSD*_*com*_), we observed that, in the cases of strong anomalous dynamics, the MSD of individual monomers *MSD*_*i*_ follows the dynamics of the center of mass in the super-Rousean regime (**Fig.6B**). Since *MSD*_*com*_ does not depend strongly on the intra-TAD strength of interaction, the transition between the small time-scale - homopolymer-like - diffusion and the large time-scale - center-of-mass-like - regime is driven by the degree of compaction of the TAD. The cross- over time between these two regimes corresponds typically to the Rouse time of the TAD (Grest and Kremer 1986), i.e. the typical time for the center of mass of the TAD to diffuse over the typical 3D physical size of the TAD (given by the average end-to-end distance 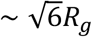) (**Fig.6B**). Overall, these results suggest that the non-monotonic - anomalous - evolution of *α*_*i*_ represents a crossover between the ‘fast’ diffusion of single monomers at short times and a ‘slow’, collective regime where loci of the same TAD move coherently at longer time-scales, as recently observed experimentally for heterochromatin compartments (Eshghi et al. 2021).

### Coherent motion of intra-TAD monomers

We then reasoned that such coherent moves might lead to correlations in the motion of monomers. We thus computed from our simulations the matrix *C*_*ij*_(Δ*t*) that describes how the displacements of monomer *i* after a time lag Δ*t* are correlated with the displacements of another monomer *j* during the same time period (see **Methods**). **Fig 7A** shows the normalized pair-correlation matrix 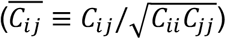 for different time-lags. 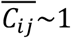 (or∼ − 1) means that, after Δ*t, i* and *j* have moved on average along the same (or opposite) direction. For the null, homopolymer model (**Fig.7A**), correlations decay uniformly towards zero as the genomic separation increases between the two loci (**Fig.7B**). If we note *s*_*corr*_ the typical genomic distance between two monomers beyond that their motions become uncorrelated 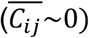, we remarked that *s*_*corr*_ augments with the time-lag from a few dozens of kbp for second-scale displacements to Mbp after hours. This is a direct consequence of the polymeric nature of chromosomes and of the conserved connectivity along the chain. For the mouse and the *Drosophila* cases, we observed patterns in the correlation matrices that are strongly related to the TAD organization as already seen by (Di Pierro et al. 2018), more compact TADs (*Drosophila*) being more impacted (**Fig.7A**) as the dynamics of intra-TAD contacts is reduced (**Suppl. Fig. S11**). This observation confirms the coherent motion of monomers inside TADs and its relation to intra-TAD compaction (**Figs.5**,**6**). For longer time-lags, correlations between monomers of the same compartment become also significant, in particular in the mouse case where compartments are more segregated (**Fig.2F-I**). Overall, this leads to larger *s*_*corr*_ values (**Fig.7B** and **Suppl. Fig. S12**), the difference with the null model increasing with the time-lag. Additionally, we looked at the spatiotemporal correlation function *C*_Δ*t*_(*r*) (**Methods**) that represents the average correlation between the displacement after a given time lag Δ*t* of two monomers initially separated by a 3D distance *r* (Liu et al. 2018) (**Fig.7C**), a quantity more easily accessible experimentally (Zidovska et al. 2013). We found that the typical size of the spatial regions with correlated motions increases with the time-lag from ∼100nm for second-scale displacement to ∼1μm after hours. These predictions remain however largely underestimated compared to experiments (few microns already after a few seconds) (Zidovska et al. 2013). We also did not find strong differences between the null model and the mouse and *Drosophila* cases, demonstrating that *C*_Δ*t*_(*r*), which is an average over all the loci, does not capture the effect of compaction on correlated motion. All this suggests that the larger spatiotemporal correlations observed experimentally are not the signature of TAD formation or A/B compartmentalization.

**Fig 7:**
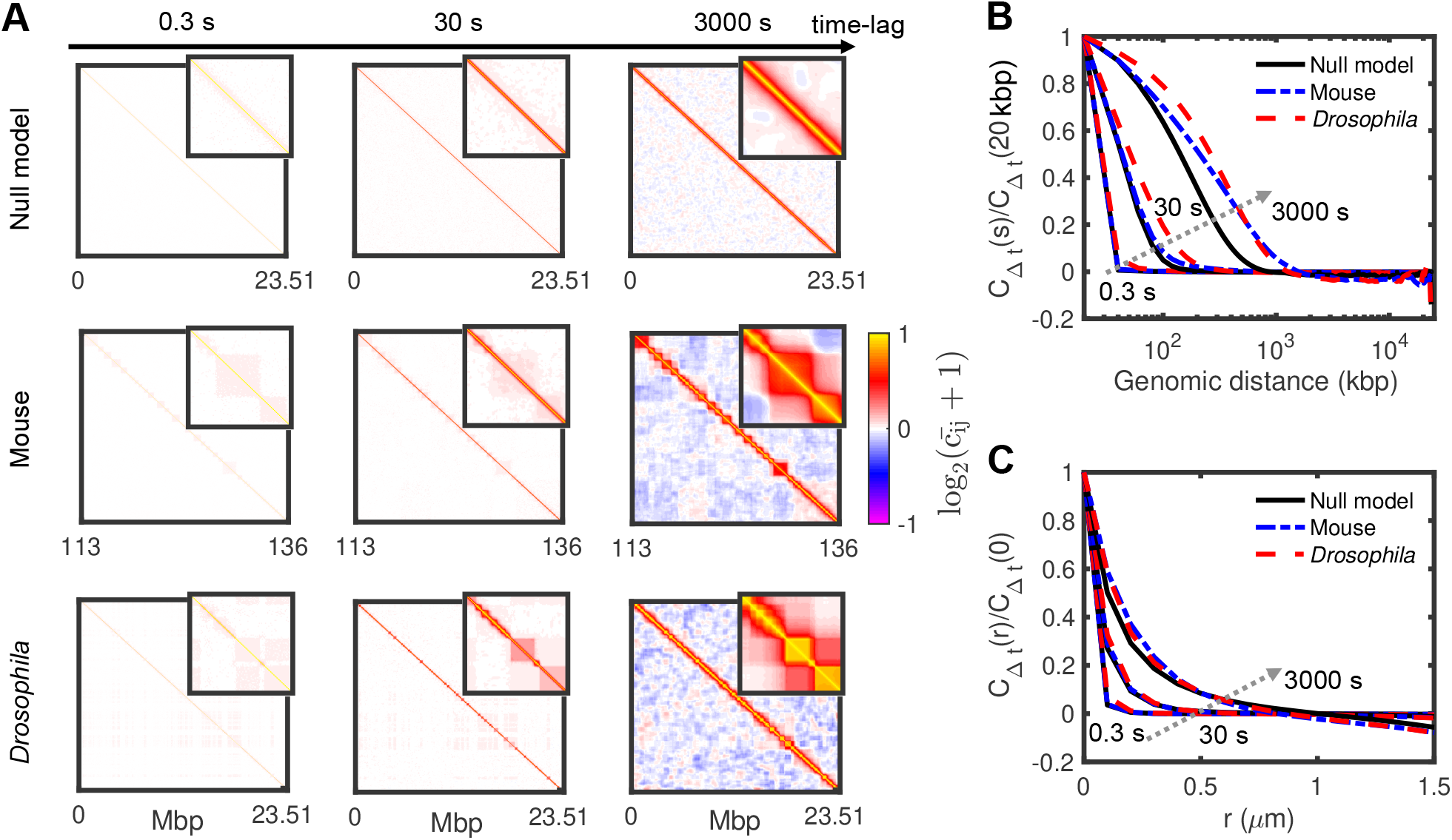
Spatiotemporal correlations of loci displacements. **(A)** Normalized matrices of pair correlation of motions 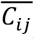 for the null model (top panels), mouse (middle panels) and *Drosophila* (bottom panels) at different time-lags. Insets represent zooms of the 2Mbpx2Mbp central parts of the matrices. **(B-C)** Normalized average correlations as a function of the genomic distance (B) and spatial distance (C) for the time-lags displayed in panel (A). Arrows indicate increasing time-lag.

### TAD formation by loop extrusion may also lead to anomalous behaviors

In our data-driven heteropolymer framework, we considered effective, passive interactions to model the patterns observed in experimental Hi-C maps. While such type of interactions, putatively driven by chromatin-binding proteins associated with specific epigenomic marks, may actually drive the formation of spatial domains like TADs in *Drosophila* (Szabo et al. 2019; Jost et al. 2014) or constitutive heterochromatin domains in higher eukaryotes (Larson et al. 2017; Strom et al. 2017), other key mechanisms like cohesin-mediated loop extrusion (Fudenberg et al. 2017) may also be involved in TAD formation. To investigate if anomalous dynamics can be observed in different mechanistic contexts, we simulated chromatin mobility for two other mechanisms on toy examples (see **Methods**).

As in our main model, the first simulation set also applies attractive interactions between loci, but only on a fraction (not all) of the monomers inside a TAD (**Fig.8A,B)**. These monomers may represent the actual binding sites of chromatin-binding proteins like PRC1 or HP1. For similar TAD compaction levels, we observed much more heterogeneous dynamics than in the uniform models seen above (**Fig.8C** and **Suppl. Fig. S13**). Monomers of the same TAD may have different MSDs: some react like in the null model while others follow anomalous behaviors, a property that seems independent of their binding site status (**Suppl. Fig. S13**). However, monomers of the same TAD still move coherently (**Fig.8D**).

The second mechanism accounts for the chromatin loop-extrusion by cohesin rings (Fudenberg et al. 2016; Sanborn et al. 2015): loop-extruding factors are loaded onto chromatin, actively extrude loops until they unbind, meet another extruder or reach TAD border (**Fig.8E**). For standard parameter values of the loop extrusion process (see **Methods**), the translocation activity of cohesin generates dynamic loops that are stabilized at TAD boundaries, leading to intra-TAD compaction (**Fig.8F**). As observed experimentally (Kakui et al. 2020), we found that the loop extrusion mechanism leads to decreased mobility compared to the null model (**Fig.8G**). We also detected more dynamical variability between monomers within a domain (**Fig.8G** and **Suppl. Fig. S14**). Strongly anomalous behaviors are found close to TAD borders where contacts mediated by the extruders are more stable, leading to collective motion of the TAD boundaries. Intra-TAD coherent motion is, instead, weaker (**Fig.8H**).

## Discussion and Conclusion

Experiments probing chromatin motion have highlighted the large heterogeneity existing inside cell nuclei and across biological conditions and have suggested that chromatin may behave sometimes like a liquid, sometimes like a gel. In this paper, we investigated chromosome dynamics using biophysical modeling in order to interpret, in a unified framework, how these different physical states and dynamical heterogeneities may emerge from the same first principles driving genome folding.

Based on a dynamical data-driven polymer model that captures the main structural properties of chromosome organization, we were able to quantify the motion of chromatin at the sub-chromosome territory level. Previous similar approaches on human chromosomes (Liu et al. 2018; Shi et al. 2018; Di Pierro et al. 2018) have shown that such types of heteropolymer models are consistent with a heterogeneous, sub-Rousean, (A/B) compartment-dependent and spatially-correlated chromatin dynamics. To go beyond these works (Liu et al. 2018; Shi et al. 2018; Di Pierro et al. 2018) and get a broader view of chromatin motion on contrasted situations, we studied in parallel 3 cases: a reference homopolymer model, and two heteropolymer models of a less compact, stem-cell-like chromatin (mESC) and a more compact, differentiated-cell-like organization (*Drosophila* Kc167).

Our results demonstrated indeed the heterogeneous locus-dependency of chromatin motion. The analysis of time-averaged MSDs computed from single-cell trajectories suggests however that great caution should be taken when interpreting the distribution of diffusion exponents from such experiments, as part of the observed heterogeneities may arise from intrinsic variabilities inherent to short trajectories. For the better defined, ensemble-averaged MSDs, we found that the observed dynamical heterogeneity reflects the various degree of compaction that may exist along the genome: while loci inside small or weakly-compacted TADs (or compartments) exhibit a rather homogeneous, fast, Rouse-like diffusion, loci inside compact TADs have a lower mobility and experience crossovers between different diffusion regimes (from Rousean to sub-Rousean to super-Rousean to Rousean modes). Using uniform models and testing several key mechanisms for TAD formation, we demonstrated that such anomalous behavior is the signature of collective, coherent motion at the level of strongly-compacted regions.

We observed that the existence of 3D chromatin domains of correlated displacements, the so-called dynamically-associated domains (DADs) (Zidovska et al. 2013; Shaban et al. 2018), emerges intrinsically from the polymeric nature of chromatin. However, the persistence of these domains is strongly related to the 3D chromosome organization: loci in the same TAD and, to a lesser extent, in the same compartment are more likely to be in the same DAD, as also observed by Di Pierro et al for human chromosomes (Di Pierro et al. 2018).

In our heteropolymer models, we integrated the multiple layers of chromatin organization using effective passive interactions, i.e. ATP-independent processes that satisfy detailed balance (Chandler 1987). It is interesting to note that using such interactions, we can capture super-Rousean regimes, not as a consequence of directed forces or active processes but as a crossover regime between a slow, coherent mode of motion at slow or intermediate time-scales to a normal Rouse-like dynamics at longer time-scales. However, our analysis reveals that this type of interactions cannot quantitatively capture the fast and large-scale average increase of correlated motion across the nucleus (Zidovska et al. 2013; Zidovska 2020), suggesting that other processes mediate this large-scale growth in spatial correlations (Liu et al. 2018), putatively via the actions of extensile motors on chromatin mediated by hydrodynamics interactions (Saintillan et al. 2018) or crosslinks and interactions with the nuclear membrane (Liu et al. 2021).

Such passive interactions might account for actual mechanisms, like polymer (micro)-phase separations (Jost et al. 2014; Falk et al. 2019; Erdel and Rippe 2018) mediated by homo- or heterotypic interactions between chromatin-binding proteins, that drive euchromatin/heterochromatin compartmentalization in many species and TAD formation in *Drosophila*. However, recently, an active process, the chromatin-loop extrusion by cohesins or condensins (Fudenberg et al. 2017; Ghosh and Jost 2020), was shown to play a central role in TAD formation in mammals (Fudenberg et al. 2016; Sanborn et al. 2015). This mechanism was suggested to act on chromatin dynamics either by slowing down chromatin motion (Kakui et al. 2020) ; or by boosting locally the mobility on a short-time scale corresponding to the loop extruder residence time at a locus (Nuebler et al. 2018). In our hands, the loop extrusion mechanism leads to an overall reduced and heterogeneous mobility. Coherent motions and anomalous behaviors are also visible for such a process but mainly for genomic regions close to TAD boundaries, where compaction is stronger and cohesin-mediated loops less dynamic.

Our work provides a unified framework to rationalize the wide variety of behaviors observed experimentally. Heterogeneity in the diffusion and exponent constants are driven by heterogeneity in TAD organization and chromatin condensation. The observed fluid-like behavior (Maeshima et al. 2016; Ashwin et al. 2020) of chromatin is likely to be associated with weakly-compacted, dynamic chromatin. This would typically correspond to stem-cell-like conditions where A and B compartments are still not entirely formed and where architectural proteins driving their organization like HP1 (Strom et al. 2017) are not fully loaded (Poonperm and Hiratani 2021). This may explain why no clear differences are globally observed between eu- and heterochromatin in U2OS (Shaban et al. 2020b), a highly plastic and transformed human cell line. The gel-like state of chromatin (Khanna et al. 2019; Eshghi et al. 2021) is associated with strongly-compacted regions which, in our examples, mainly correspond to TADs but would also be observed for any chromatin structure with similar degrees of compaction. Within our framework, this corresponds to the weak gelation of a polymeric system (de Gennes 1979; Douglas 2018) where changes in the effective internal friction or viscosity (Poirier and Marko 2002; Soranno et al. 2012; Socol et al. 2019) emerges from reversible crosslinks (**Suppl. Fig. S15**). The dynamic signature of such a gel-like state relies on its low mobility, the presence of local coherent motion and anomalous behaviors with slow internal dynamics and crossovers between different diffusion regimes for individual loci (see above). By combining live-imaging and modeling, Khanna et al (Khanna et al. 2019) have described such a gel state at the immunoglobulin heavy chain locus in pro-B cells, a highly compacted region. They observed very slow, sub-Rousean, internal motion between V and DJ segments and a crossover between super-Rousean and Rousean regime for individual MSD of the DJ segment, which is consistent with our analysis of strongly self-interacting regions **(Suppl. Fig. S16**). At the nuclear scale, this gel state could be only localized on a few compacted regions like on centromeres or telomeres, even in plastic cells (Bronshtein et al. 2015; Eshghi et al. 2021). For example, recent experiments by Eshghi et al (Eshghi et al. 2021) showed that, in mESCs, loci within dense heterochromatin compartments exhibit anomalous, gel-like MSDs with a crossover towards collective motion for the entire compartment. We expect the gel-like dynamics to become more and more predominant as cell differentiation progresses and compartments like heterochromatin achieve their final compaction state (Lerner et al. 2020). This may lead in extreme cases to a solid state with an almost arrested dynamics of chromatin as recently suggested by FRAP analysis on eu- and heterochromatin regions of mouse embryonic fibroblast cells (Strickfaden et al. 2020).

Are the different dynamical regimes of chromatin only readouts of the mechanisms driving chromosome organization or do they carry, in addition, specific biological functions? Regulating chromatin mobility may directly impact gene transcription (Maeshima et al. 2020) by controlling the timing of contact between promoters and distant enhancers and thus the gene bursting frequency (Bartman et al. 2016; Chen et al. 2018). Similarly, the regulation of the epigenomic landscape may be affected by the local dynamical regimes as the kinetics of spreading and maintenance of an epigenomic signal would depend on the capacity of the genomic loci where histone-modifying enzymes are bound to, to explore more or less rapidly their 3D neighborhood and thus to allow the propagation of the corresponding histone modification (Jost and Vaillant 2018; Oksuz et al. 2018). For example, active marks usually found in less-compact, more-fluid compartments have a faster dynamics of maintenance than inactive marks (Alabert et al. 2015). More generally, we expect the regulation of a genomic region to be more sensitive and to adapt more efficiently to variations in a fluid-like environment than in a gel-like environment that, on contrary, may protect it from spurious, non-persistent perturbations.

A better characterization of the regulation of the fluid-, gel- or solid-like states of chromatin motion would require the development of upgraded polymer models integrating the main passive and active processes driving genome folding but also the development of new experimental approaches allowing to quantify simultaneously the dynamics of many loci (whose genomic positions are known) at high spatial and temporal resolution. Recent progresses in multiple loci (Zhou et al. 2017) and in super-resolution (Alexander et al. 2019; Chen et al. 2018; Brandão et al. 2020; Barth et al. 2020) live imaging would make it possible to test quantitatively the relations between mobility, coherent motion and compaction predicted by our polymer models and the role of basic mechanisms such as phase separation or loop extrusion in regulating chromosome dynamics. For example, live-imaging the coupled structural and dynamical responses of chromatin after rapid *in vivo* perturbations of the amount of key chromatin-binding complexes using the auxin-inducible-degron system (Schwarzer et al. 2017; Dobrinić et al. 2021) or after modifications of their self-interacting capacities using optogenetics (Shin et al. 2019; Shimobayashi et al. 2021) would allow to directly investigate how chromatin dynamics respond to changes in chromatin compaction.

## Methods

### Analysis of Hi-C data

The experimental Hi-C datasets were downloaded from the sequence read archive (SRA) (**Table 1**) using *fastq-dump* (https://github.com/ncbi/sra-tools/wiki). Each experiment has been processed (i.e. FASTQ quality checks, Mapping to reference genomes, Filtering to remove non-informatic reads, and Merging data together) through the TADbit pipeline (https://github.com/3DGenomes/TADbit) (Serra et al. 2017). Then, we normalized data using the Vanilla method at 10 kbp resolution (Lieberman-Aiden et al. 2009). After generating the Hi-C map, we employed the IC-Finder tool to find TAD boundaries (Haddad et al. 2017). The A/B compartments have been identified by the *hic_data*.*find_compartments* tool from the TADbit suite, using Principal Component Analysis (PCA) of the observed/expected matrices. Finally, we inferred chromatin loops by using *hicDetectLoops* tool from the HiCExplorer package (https://github.com/deeptools/HiCExplorer) (Wolff et al. 2020).

### Polymer models and simulations

Each genomic segment under investigation is modeled as a semi-flexible self-avoiding polymer in which one monomer consists of 2 kbp of genome and has a diameter of 50 nm (**Fig 2A**). The chain is moving on a FCC lattice under periodic boundary conditions to account for confinement by other genomic regions, as described in (Ghosh and Jost 2018).

The Hamiltonian of a given configuration is given by

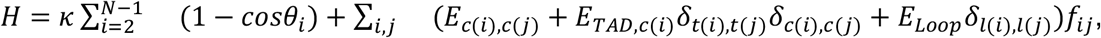

with *f*_*ij*_=1 if monomers *i* and *j* occupy nearest neighbor sites on the lattice (= 0 otherwise). The first term in *H* accounts for the bending rigidity of the chain with *κ* the bending stiffness and *θ*_*i*_ the bending angle between monomers *i* − 1, *i* and *i* + 1. The second term refers to contact interactions driven by the compartment (*c*(*i*)), TAD (*t*(*i*)) or loop (*l*(*i*)) state of each monomer. *E*_*c*(*i*),*c*(*j*)_ (either *E*_*AA*_, *E*_*BB*_ or *E*_*AB*_) stands for compartment-compartment interactions, *E*_*TAD,c*(*i*)_ (either *E*_*TAD,A*_ or *E*_*TAD,B*_) for intra-TAD interactions, *E*_*Loop*_ for loop interactions and *δ*_*m,n*_ = 1 if *m* = *n* (=0 otherwise). In the homopolymer model, all interaction parameters except *κ* were set to zero.

The lattice volumic fraction (∼0.5) and bending energy (1.2 kT) were chosen to simulate the coarse-grained dynamics of a chromatin fiber of Kuhn length ∼100 nm (Socol et al. 2019; Arbona et al. 2017) and of bp-density ∼0.01 bp/nm^3^, typical of mammalian and fly nuclei (Milo et al. 2009). Dynamics of the chain was simulated using kinetic Monte-Carlo, starting from unknotted initial configurations, as detailed in (Ghosh and Jost 2018). Each Monte-Carlo step (MCS) consists of N (total number of monomers) local moves including reptation moves. Such implementation has been shown to well reproduce the structural and dynamical properties of long, confined polymers (Olarte-Plata et al. 2016; Ghosh and Jost 2018). For each simulation, we discarded the first 10^6^ MCS to reach steady state and then stored snapshots every 10^2^ MCS during 10^7^ MCS. For each situation (homopolymer, mouse and fly heteropolymer models), 20 different independent trajectories were simulated starting from different initial configurations.

### Parameter inference of the heteropolymer models

To infer parameters in both (mouse and *Drosophila*) cases, we first mapped TADs, compartments and loops extracted from the experimental Hi-C map into the heteropolymer model and then iteratively adjusted the interaction parameters to optimize the correspondence between the predicted and experimental Hi-C maps. For simulations, we estimated the contact frequency *P*_*ij*_(*sim*) between any pair *(i,j)* of monomers as the probability to observe *i* and *j* at a distance less than a cutoff value *r*_*c*_ in all our snapshots. To quantitatively compare simulated contact probabilities to experimental Hi-C map, we transformed the experimental contact frequencies (*H*_*ij*_) to contact probabilities (*P*_*ij*_(*exp*)) using the following relation 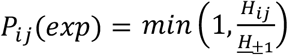, where *H*_+1_ is the median value of {*H* _*i,i* +1_} (Szabo et al. 2018). Our criteria for finding the optimal parameter values (energies *E*_*AA*_, *E*_*BB*_, *E*_*AB*_, *E*_*TAD,A*_, *E*_*TAD,B*_, *E*_*loop*_ and cutoff distance *r*_*c*_) were the maximization of the Pearson’s correlation between simulated and experimental matrices and the minimization of the χ^2^ score, 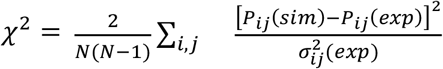, where 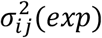 is the experimental standard deviation on *P*_*ij*_(*exp*) estimated over experimental replicates. The optimization was done using a grid-search algorithm over the parameter values in the range [-0.3:0] kT for energy parameters and in the range of [70:200] nm for *r*_*c*_. In **Table 2** we summarized the optimal values for each case with their corresponding χ^2^ and Pearson’s correlation. For comparison, we also computed the correlation between *P*_*ij*_(*exp*) and random permutations of the *P*_*ij*_(*sim*) matrix where bins corresponding to the same inter-loci genomic distance |*j* − *i*| were randomly shuffled in order to preserve the average contact frequency decay as a function of the genomic distance. For these random matrices, we found a correlation of 0.8414 ± 0.0001 for the mouse case and 0.7852 ± 0.0008 for the *Drosophila* case (compare to the optimal values in Table 2), showing that our models predict much more than just the average behavior.

### Time mapping of simulations

In kinetic Monte-Carlo (KMC) frameworks, the dynamics of particles depend on the acceptance ratio of local trial moves. For very small step size in trial moves and very weak potentials, KMC results are equivalent with Brownian dynamics, while for longer step sizes and stronger interactions, the KMC time steps need to be rescaled (Sanz and Marenduzzo 2010 ; Bal and Neyts 2014). Since we are simulating a unique chain in the box in absence of external forces, we expect the MSD (*g*_3_) of the center of mass of the whole polymer to be independent of the investigated model (homopolymer or heteropolymer). However, we found (**Suppl. Fig. S17A**) *g*_3_ ≅ 5.70 × 10^−8^(μ*m*^2^/*MCS*)Δ*t* for the homopolymer model, *g*_3_ ≅ 5.50 × 10^−8^(μ*m*^2^/*MCS*)Δ*t* for the mouse case and *g*_3_ ≅ 2.85 × 10^−8^(μ*m*^2^/*MCS*)Δ*t* for the *Drosophila* case, where *g*_3_ is measured in μ*m*^2^ and Δ*t* in *MCS*. Therefore, we rescaled time *MCS* → *MCS*^*^ for each model to have similar *g*_3_ for all cases (**Suppl. Fig. S17B**). The rescaled time for mouse is *MCS*^*^ = 0.96 *MCS* and for *Drosophila MCS*^*^ = 0.50 *MCS*. As expected, the rescaled time is dependent on the strength of interaction, in a way that *MCS*^*^ decreases for stronger interactions. Then, to translate *MCS*^*^to real time (sec), we compare the average simulated 2D-MSD of single loci (in μ*m*^2^), to the typical value of 0.01(μ*m*^2^/*sec*^0.5^) × Δ*t*^0.5^(with Δ*t* in sec) that has been measured in yeast (Hajjoul et al. 2013) and that corresponds to an average MSD also observed in other species (**Fig.1A**). From this time mapping, we found that each *MCS*^*^corresponds to ∼3 msec of real time.

### Uniform heteropolymer and loop extrusion models

In order to separately investigate the effects of TAD compaction, TAD arrangements and compartments, we introduced toy polymer models. In the first step, in order to explore the effects of TADs (i.e. TAD length and intra-TAD interaction), we considered “uniform” models where 20Mbp-long chromosomal segments are partitioned into consecutive TADs of uniform lengths (**Suppl. Fig. S5**-**S7**). Then, to investigate the role of TADs arrangement, we partitioned the polymer into TADs alternating between two different lengths (**Suppl. Fig. S8** upper panels, **Suppl. Fig. S9**). Finally, to investigate the effect of compartments, we alternatively assigned A and B compartments to the uniform TAD models with *E*_*BB*_ *<* 0 and *E*_*AA*_ = *E*_*AB*_ = 0 (**Suppl. Fig. S8** bottom panels, **Sup Fig. S10**).

In another scenario, we introduced toy models to investigate the effect of discrete binding sites on chromosome dynamics. Similar to the uniform models described above, we partitioned the chromosome into TADs of uniform length. In each TAD, we assigned randomly, to a fixed proportion *ρ* of monomers, the so-called ‘binder’ monomers, the capacity to interact with an attractive energy *E*_*b*_ with other ‘binders’ of the same TAD (**Fig.8A**). Other monomers are considered as neutral monomers. To have a similar effective intra-TAD interaction and consequently similar compaction levels, we adjusted *E*_*b*_ and *ρ* to satisfy *ρ*^2^*E*_*b*_ = *E*. In **Fig.8** and **Suppl. Fig. S13**, we chose *E* = −0.05 *kT*.

We also investigated the effect of loop-extrusion activity on chromosome dynamics (**Fig. 8E**). We considered *N*_*tot*_ loop extruders that may bind (with rate *k*_*b*_) or unbind (with rate *k*_*u*_) from the polymer. Bound extruders are composed of two legs that walk on opposite directions along the polymer at a rate *k*_*m*_, these legs are linked together by a spring of energy *E*_*a*_Δ*r*^2^ with *E*_*a*_ = 10*kT* and Δ*r* the 3D distance on the lattice between the two legs. If one leg is on a monomer corresponding to a TAD border, its moving rate is set to zero, assuming that this border is enriched in CTCF binding sites (Rao et al. 2014; Dowen et al. 2014) that are known to stop or limit the loop extrusion of cohesins (Nora et al. 2020, 2017). We assumed that two extruding legs walking in opposite directions cannot cross.

**Fig 8:**
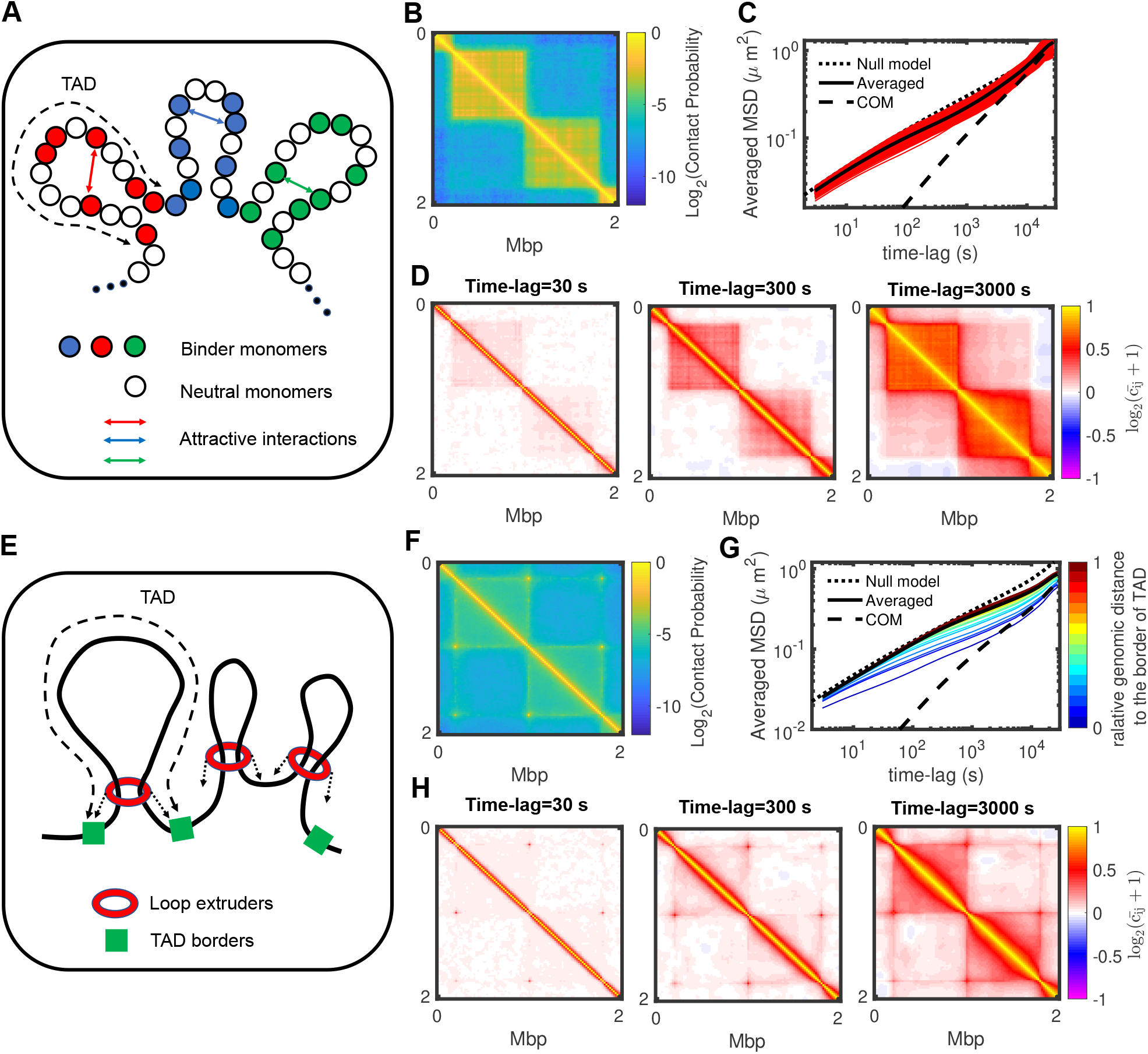
Effects of alternative mechanisms for TAD formation on anomalous behaviors. **(A)** Schematic representation of a binder model for TAD formation. **(B)** 2Mb zoom of the predicted Hi-C map for a toy model with 800kbp-long TADs, a density of binder monomers of 0.5 and an attractive interaction of -0.2 kT (see **Suppl. Fig. S13** for other examples). **(C)** Ensemble-averaged MSDs of the binder monomers with their average and the MSD of the TAD center of mass for the same parameters as in (B). **(D)** 2 Mb zooms in the matrix of pair correlations 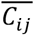 for different time-lags for the example shown in (B). **(E)** Schematic representation of the loop-extrusion model. **(F)** 2 Mb zoom of the predicted Hi-C map for a toy model with 800kbp-long TADs and about 80 bound extruding factors (see **Suppl. Fig. S14** for other examples). **(G)** Ensemble-averaged MSDs colored by the relative position of the monomer to the nearest TAD border for the same parameters as in (F). **(H)** 2 Mb zooms in the matrix of pair correlations 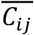 for different time-lags for the example shown in (F).

In our kinetic Monte-Carlo framework, in addition to the standard trial moves of the monomers based on the Hamiltonian of the system (see above) complemented with the spring-like interactions, at every MCS, *N*_*tot*_ trial attempts to bind or unbind extruders and 2*N*_*tot*_ trial attempts to move a leg of a bound extruder are also performed. *k*_*u*_ was fixed to fit the experimentally-observed life time of bound cohesin on chromatin (∼20 min) (Hansen et al. 2017), *k*_*b*_ such that the proportion of bound extruders *k*_*b*_/(*k*_*b*_ + *k*_*u*_) is about 40% (Cattoglio et al. 2019) and *k*_*m*_ such that the extruding speed rate matched *in vitro* experimental values (∼50*kbp*/*min*) (Golfier et al. 2020). In **Fig.8** and **Suppl. Fig. S14**, we explored the effect of TAD length and *N*_*tot*_on structure and dynamics. For example, *N*_*tot*_ = 200 implies ∼80 bound extruders in average acting on the chain, corresponding to a density of ∼1 extruder every 250 kbp.

### Fitting of the diffusion exponent and constant

To extract the diffusion constant, *D*_*i,c*_, and exponent, *α*_*i,c*_ for each trajectory *c* and each monomer *i*, we fitted the time-averaged mean squared displacement *MSD*_*i,c*_ by a power-law 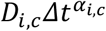 using the MATLAB function *polyfit(LogMSD*_*i,c*_,*Log*Δ*t*, *1)* over more than three decades. Note that the unit of *D*_*i,c*_ depends on *α*_*i,c*_ and is 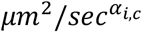 (see **Suppl. Fig. S2**,**3**). For the ensemble-averaged mean squared displacement *MSD*_*i*_, we assumed that the diffusion exponent is a function of the time-lag 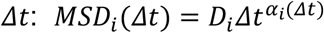 or, in logarithmic scale *Log MSD*_*i*_ = *Log D*_*i*_ + *α*_*i*_*Log*Δ*t. α*_*i*_(Δ*t*) is then given by the local slope of the log-log MSD curves:

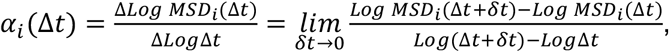

and *D*_*i*_ by

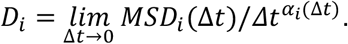

Note that, since *α*_*i*_(Δ*t* → 0)∼0.5, the unit of *D*_*i*_ is μ*m*^2^/*sec*^0.5^.

### Two-dimensional density plot

To construct the 2D density plot of chromosomes shown in **Fig.2F,H**, the intensity at a position (*x, y*) is given by the sum over all monomers of a 2D Gaussian function mimicking the point-spread-function (PSF) of a microscope:

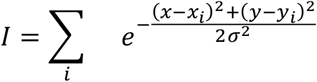

with (*x*_*i*_, *y*_*i*_)the position of monomer *i* and *σ* = 300nm chosen to get a spatial resolution of 100 nm in the (*x, y*)-plane.

### Spatiotemporal correlation function

We defined *C*_*ij*_(Δ*t*), the pair-correlation of the displacement vectors after a time-lag Δ*t* of the monomers *i*-th and *j*-th, as (Di Pierro et al. 2018)

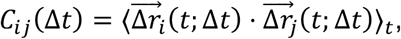

with,

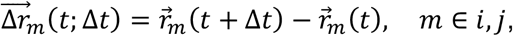

where 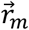 is the position of monomer *m* with respect to the center of mass of the chain. Then, we calculated the averaged correlation *C*_Δ*t*_(*s*) as a function of the genomic distance *s* between monomers by averaging over all *C*_*ij*_ with same genomic distance:

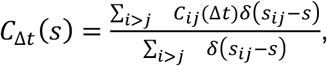

where, *s*_*ij*_ is the genomic distance between monomers *i*-th and *j*-th, and *δ*(*x*) is the Kronecker delta function. Additionally, we defined the spatiotemporal correlation function (i.e., averaged correlation as a function of spatial distance), *C*_Δ*t*_(*r*), as (Liu et al. 2018)

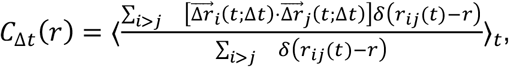

where, *r*_*ij*_(*t*) is the spatial distance between monomers *i*-th and *j*-th.

Note that if we did not correct for the movement of the center of mass (**Suppl. Fig. S18**) we observed very long-range correlations between loci as the time-lag increases, which reflects the global motion of the center of mass.

## Software availability

Codes used to simulate the heteropolymer and the loop extrusion models are available as on https://github.com/physical-biology-of-chromatin/LiquidGel_ChromSimu and in the Supplemental Code file.

## Competing interests

The authors declare no competing interests.

## Acknowledgments

We are grateful to Cédric Vaillant, Geneviève Fourel, Haitham Shaban, Kerstin Bystricky and the members of the Jost lab for fruitful discussions. DJ acknowledges Agence Nationale de la Recherche [ANR-18-CE12-0006-03, ANR-18-CE45-0022-01] for funding. We thank the PSMN (Pôle Scientifique de Modélisation Numérique) of the ENS de Lyon for computing resources. MDS acknowledges a STSM Grant from COST Action CA17139 supported by COST (European Cooperation in Science and Technology).

## Author contributions

HS and DJ designed the research, HS and DJ performed the research, HS, MDS and DJ analyzed the data, HS and DJ wrote the paper with significant inputs from MDS.

## Notes

### Competing Interest Statement

The authors have declared no competing interest.

