## Supplemental file for "Spatial organization of chromosomes leads to heterogeneous chromatin motion and drives the liquid- or gel-like dynamical behavior of chromatin"

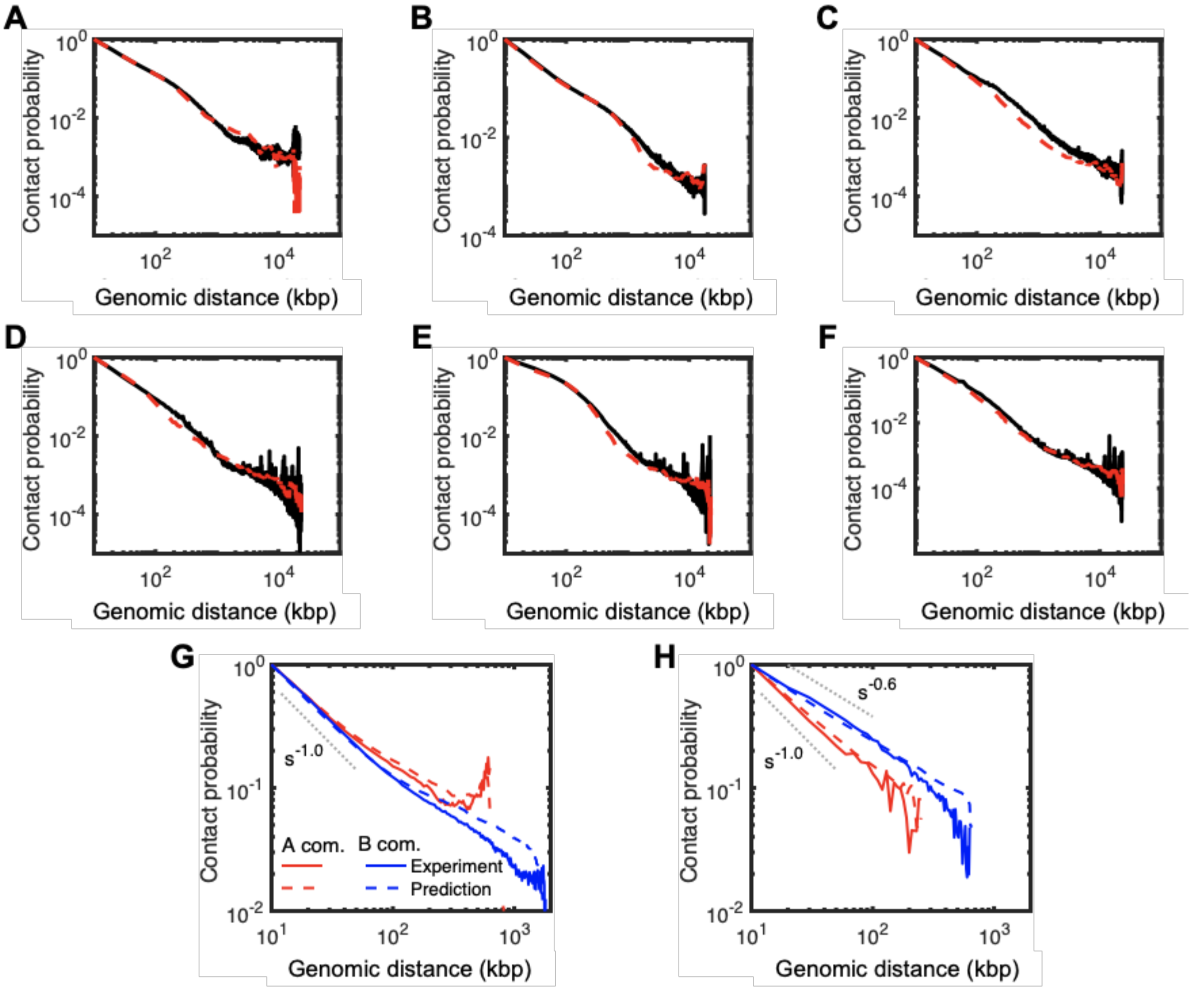


**Suppl. Fig. S1: Comparison between contact probabilities. (A-F)** Average contact probabilities between two monomers distant by a given genomic distance and localized both in the A compartment (A,D), both in the B compartment (B,E) and one in A and one in B (C,F), for the mouse (A-C) and the *Drosophila* (D-F) cases. Full black lines correspond to experimental data, red dashed lines to heteropolymer predictions. **(G,H)** Average intra-TAD contact probabilities as a function of the genomic distance for monomers in A (red) or B (blue) compartments for the mouse (G) and *Drosophila* (H) cases. Full lines correspond to experimental data, dashed lines to heteropolymer predictions. Repressive/B compartment in *Drosophila* Kc167 is more compact than active/A compartment, while they exhibit similar compaction for mouse ESCs. The heteropolymer model is able to quantitatively describe these structural properties for both examples.

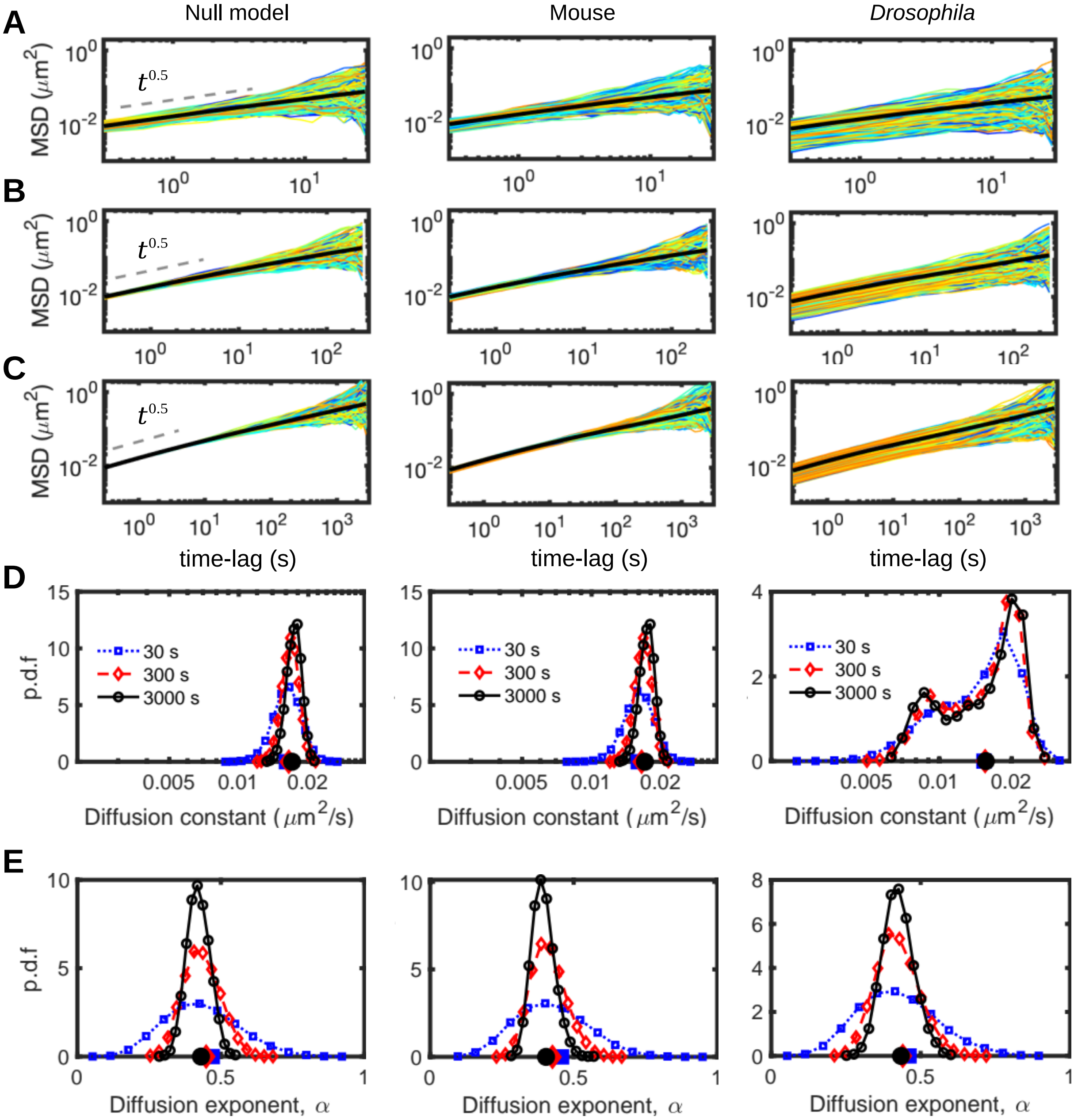


**Suppl. Fig. S2: Effects of trajectory length on the measure of dynamical indicators.** Time-averaged $MSD_{i,j}$ of single trajectories (different colors) for trajectories of length 30 s **(A)**, 300 s **(B)** and 3000 s **(C)**. The black, full curves are the averaged MSD over all trajectories. The distributions of diffusion exponent **(D)** and exponent **(E)** of the MSD shown in (A), (B) and (C) panels. Their averaged values are shown on the horizontal axes. Increasing the trajectory length will reduce the statistical heterogeneity and the contribution of the different structural layers on local mobilities is clearer.

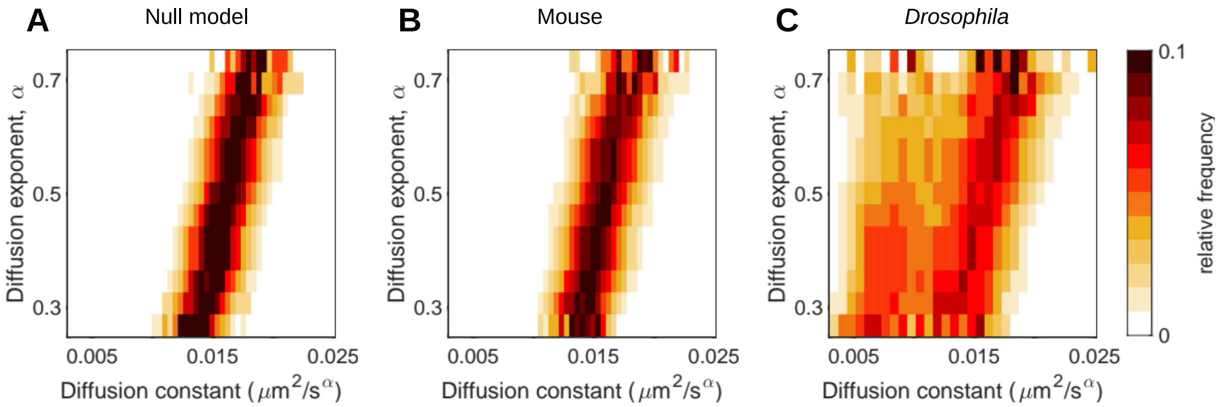


**Suppl. Fig. S3: Distributions of diffusion exponent and constant obtained from time-averaged MSDs of 30s-long trajectories. (A-C)** Relative frequency of diffusion constant for a given diffusion exponent in the null (A), mouse (B) and *Drosophila* (C) cases. For each case, we first computed the joint histogram of (diffusion constant, diffusion exponent) with a binning of $\Delta\alpha = 0.05$ for the diffusion exponent. Then for each $\alpha$-bin, we normalized the corresponding distribution of diffusion constant values. Null and mouse models behave similarly with, for each diffusion exponent regime, a monomodal distribution for the diffusion constant. The *Drosophila* case exhibits bimodal distributions, with one mode as in the null and mouse models and another mode characterized by a slower dynamics.

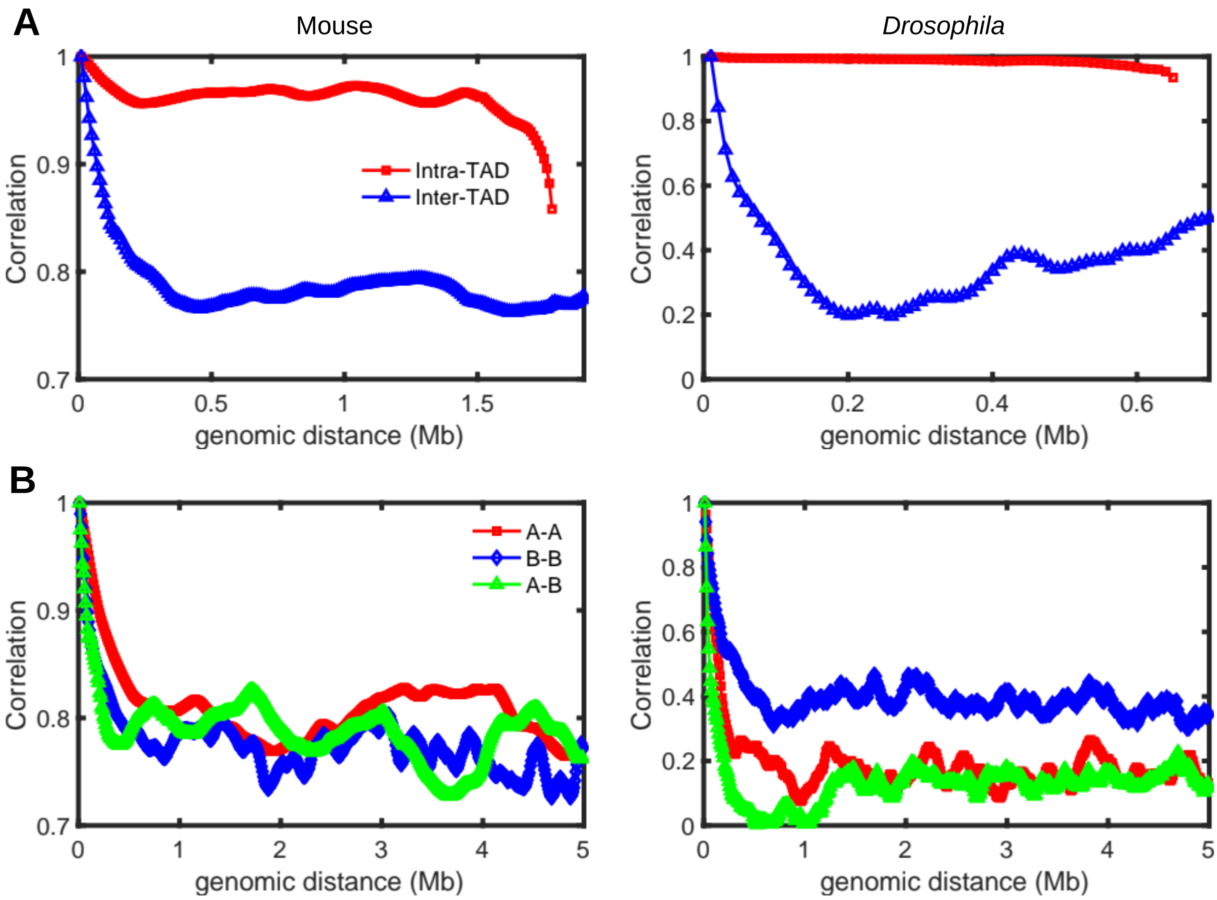


**Suppl. Fig. S4: Coupling between organizational layers and dynamics.** For every pair of loci (*i,j*), we computed the Pearson correlation between the time evolutions $\alpha_i(\Delta t)$ and $\alpha_j(\Delta t)$ of their diffusion exponents (see **Fig.4A-C** of the main text). **(A)** Average correlation values for two loci inside the same TAD (red line) or in different TADs (blue lines) as a function of their genomic distance for the mouse (left) and *Drosophila* (right) cases. In both cases, time-evolution of $\alpha_i$ are strongly correlated (>0.95) between intra-TAD monomers compared to inter-TAD monomers. This is particularly visible in the *Drosophila* case where intra-TAD compaction and anomalousness are enhanced. **(B)** As in (A) but for two loci being in the same compartments (red for A, blue for B) or in different compartments (green). While monomers of the same compartment are not significantly more correlated in mice (where compartmentalization is mild), monomers in the more compact, B compartment in *Drosophila* tend to be more correlated (~0.4) than in A or inter-compartment.

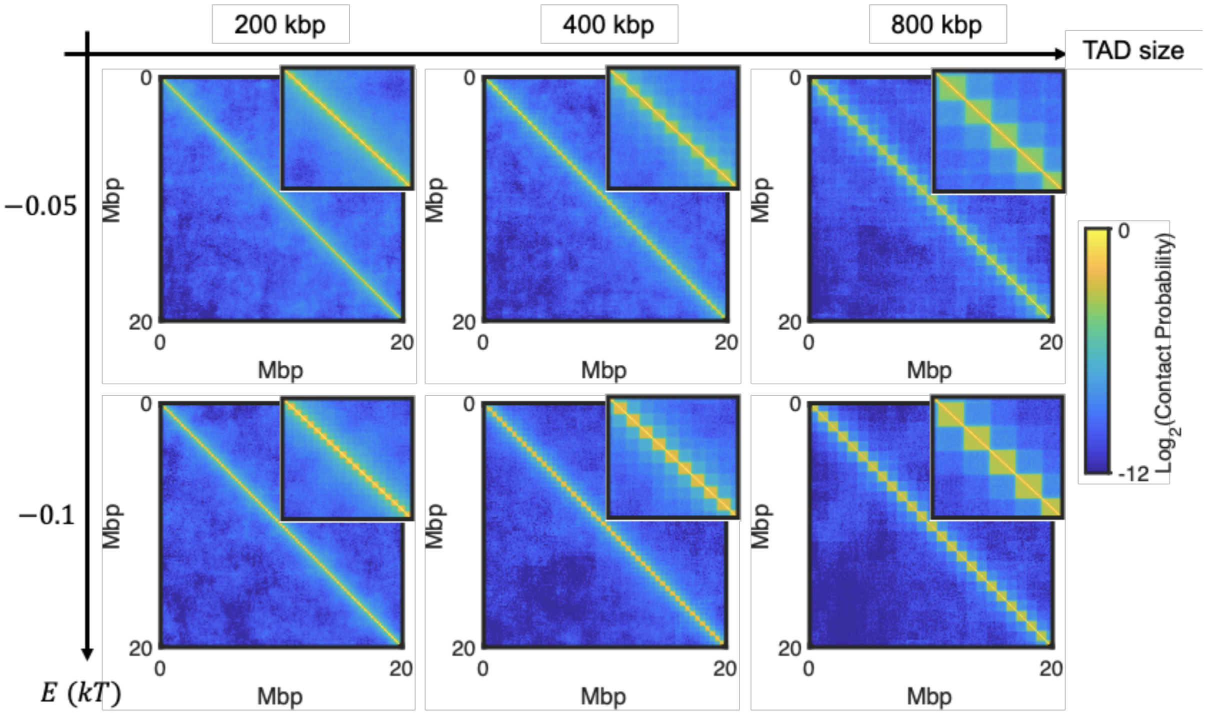


**Suppl. Fig. S5: Hi-C maps of uniform models with different TAD lengths and strengths of interaction.** Insets show 4 Mbp zoom views of the matrices. Increasing the TAD length and intra-TAD interaction leads to higher intra-TAD contact probabilities.

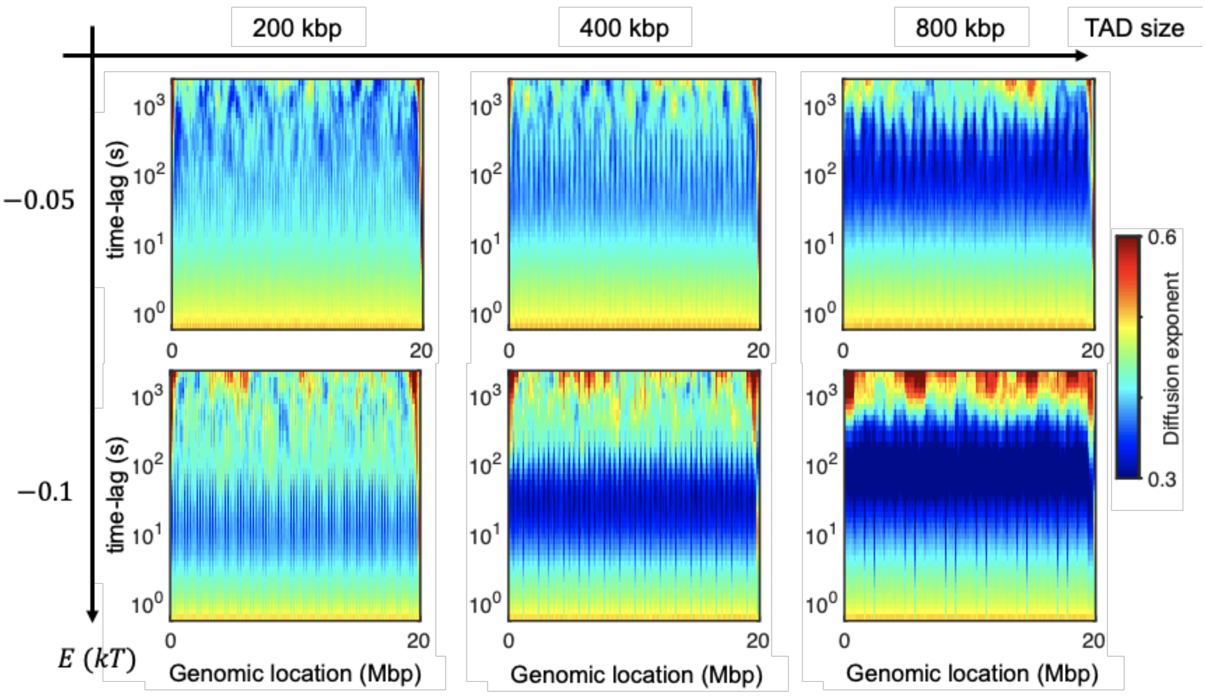


**Suppl. Fig. S6: Time-evolution of diffusion exponents of all monomers in uniform models for different TAD lengths and strengths of interaction.** Loci in more compact regions exhibit steeper anomalous behaviour.

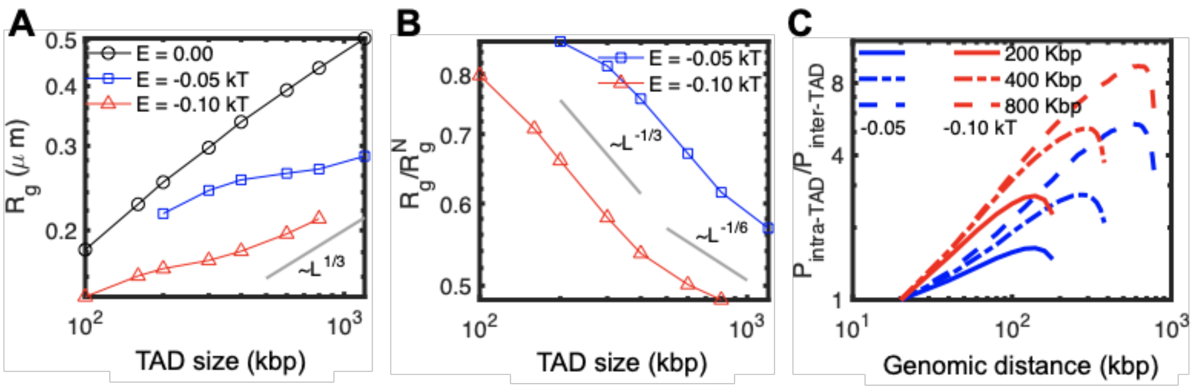


**Suppl. Fig. S7: Structural properties and compaction level of uniform models.** Radius of gyration **(A)** and ratio of $R_g/R_g^N$ **(B)** as a function of the TAD length for different strengths of interaction. **(C)** The ratio of intra-TAD to inter-TAD contact probabilities ($P_{intra-TAD}/P_{inter-TAD}$) as a function of the genomic distance for different TAD length and two different strengths of interaction. The compaction level of TADs increases by TAD length and intra-TAD interaction, in which we obtain a transition from coil state at short TAD lengths ($R_g \sim L^{1/2}$) to crumpled globule structure at large TAD lengths ($R_g \sim L^{1/3}$).

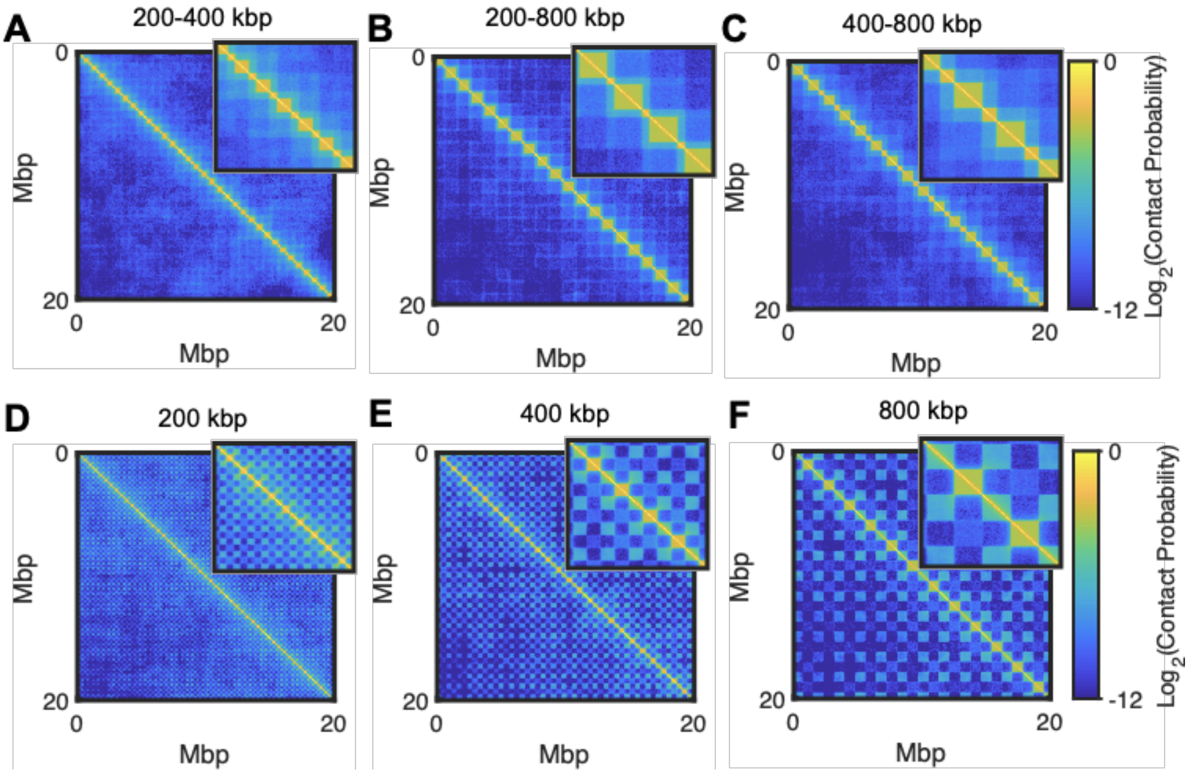


**Suppl. Fig. S8: Hi-C maps of different variations of the uniform models.** **(A-C**) Predicted Hi-C maps for models with TADs lengths alternating between two values: $(L_1, L_2) = (200{,}400)$ (A), $(200{,}800)$ (B) and $(400{,}800)$ kbp (C). **(D-F)** Predicted Hi-C maps for models with uniform TAD length (200 kbp (D), 400 kbp (E) and 800 kbp (F)) with compartment interactions. Insets show a 4x4 Mbp zoom into matrices. Bigger TADs increase the compaction level of neighboring shorter TADs**.**

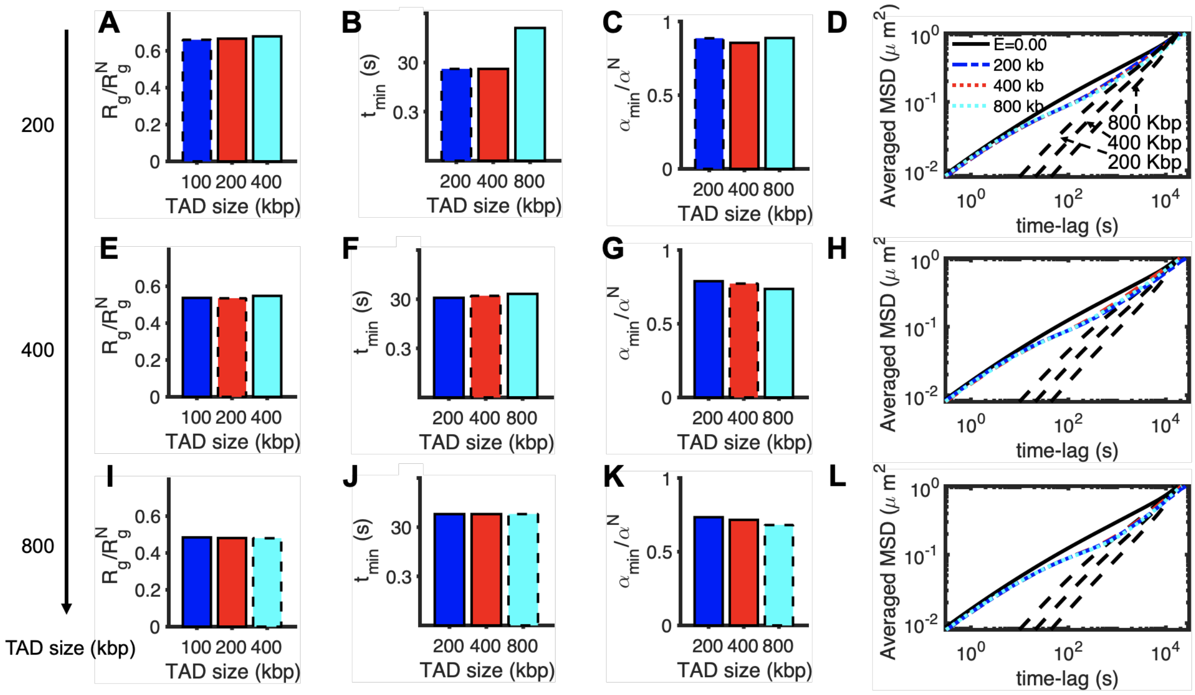


**Suppl. Fig. S9: Effect of neighboring domains on local dynamics.** Comparison of $R_g/R_g^N$ **(A)**, $t_{min}$ **(B)** and $\alpha_{min}/\alpha^N$ **(C)** for a domain of length 200 kbp having neighboring domains of different lengths, 200 kbp (blue bars), 400 kbp (red bars) and 800 kbp (cyan bars). Bars with dashed lines indicate the uniform cases. **(D)** MSD of TADs of size 200 kbp with neighboring TADs of 200 kbp (blue, dash-dotted curve), 400 kbp (red, dashed curve) and 800 kbp (cyan, dotted curve). MSDs of TADs in the homopolymer model (black, full curve) and MSDs of the center of mass of TADs of size 200, 400 and 800 kbp (black, dashed curves) are shown. **(E)**, **(F)**, **(G)** and **(H)** as in (A), (B), (C) and (D) but for a 400 kbp TAD, and **(I)**, **(J)**, **(K)** and **(L)** for a 800 kbp TAD. Bigger TADs can alter the compaction and anomalous behaviour of neighboring smaller TADs.

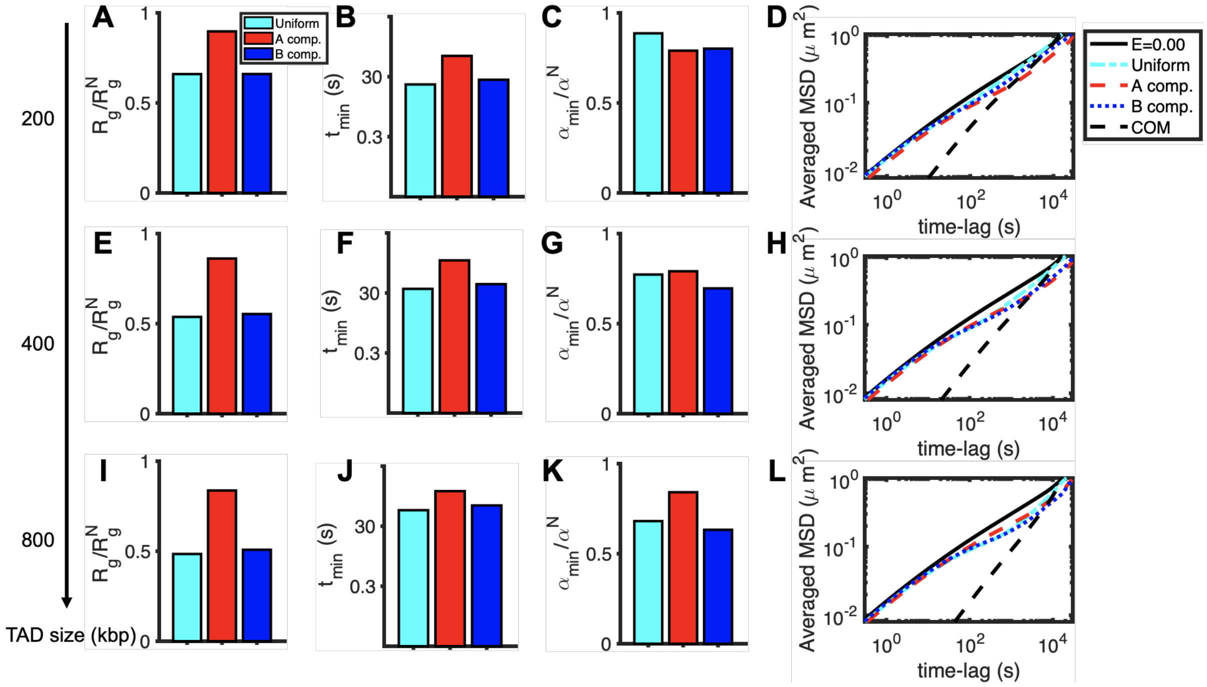


**Suppl. Fig. S10: Effects of compartmentalization on local dynamics.** Comparison between the simple uniform case (cyan bars) and the A (red bars) and B (blue bars) compartments in the uniform compartment models (see text and Suppl. Fig S11D-F) of $R_g/R_g^N$ **(A)**, $t_{min}$ **(B)** and $\alpha_{min}/\alpha^N$ **(C)** for a TAD of 200 kbp and $E = -0.1\ kT$. **(D)** Comparison of MSD of TADs in the homopolymer model (black, full curves), simple uniform model (cyan, dash-dotted curves) and the A (red, dashed curves) and B (blue dotted curves) compartments in the uniform compartment model for domain length 200 kbp and $E = -0.1\ kT$. The averaged MSDs of the center of mass of TADs are shown by black dashed curves. **(E)**, **(F)**, **(G)** and **(H)** as in (A), (B), (C) and (D) but for 400 kbp TAD, and **(I)**, **(J)**, **(K)** and **(L)** for 800 kbp TAD.

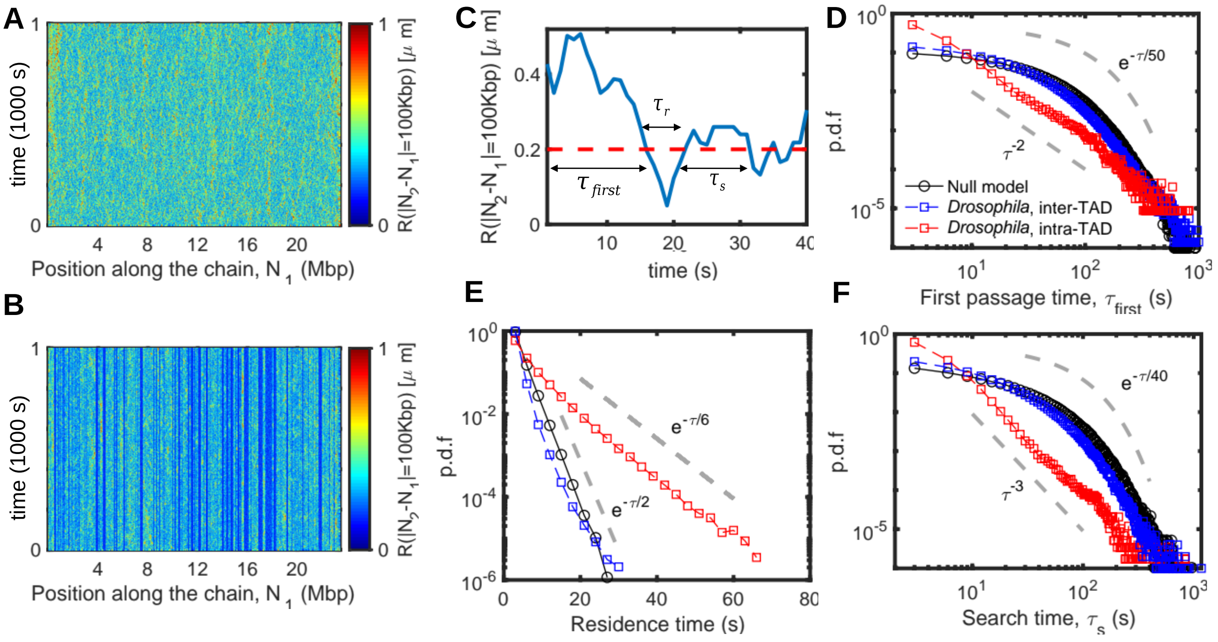


**Suppl. Fig S11: Intra-TAD slow dynamics. (A,B)** Time-evolution of spatial distances between all pairs of monomers separated by 100 kbp along the genome for the homopolymer model **(A)** and the *Drosophila* case **(B)**. The horizontal axes represent the position of the first monomer $N_1$ along the polymer. Intra-TAD relative distances are less fluctuating than inter-TAD distances. **(C)** Typical time-evolution of the distance between two monomers separated by 100kbp. The arrows illustrate the first passage time $\tau_{first}$, the residence time $\tau_r$, and search time $\tau_s$. The red dashed line represents the cut-off distance $d_c = 200\ nm$. **(D-F)** Probability distribution functions (p.d.f) for $\tau_{first}$ (B), $\tau_r$ (C) and $\tau_s$ (D) in the homopolymer case (black) and for Drosophila intra-TAD (red) and inter-TAD (blue) pairs of monomers. The grey dashed lines represent power-law or exponential fit. Note that in panels B and D, p.d.fs are shown in a log-log plot while in panel C it is in a log-lin plot. For $\tau_{first}$, intra-TAD pairs tend to meet more rapidly (average ~10 s) than inter-TAD and null model pairs (average ~50 s) (Fig.8B). The distribution of intra-TAD $\tau_{first}$ exhibits a scaling law while inter-TAD and null model pairs distribution is exponential. Residence times ($\tau_r$) are exponentially distributed for all pairs with a decay time 3-fold larger for intra-TAD pairs (~6 s for intra-TAD vs ~2 s for inter-TAD and null model pairs) suggesting stable contacts and a slow relative dynamics between intra-TAD monomers. The distributions of $\tau_s$ are almost identical for inter-TAD and null model pairs with an exponential decay characterized by long waiting times (~40 s). Intra-TAD search times are much smaller (average value ~ 6 s) and characterized by a power-law.

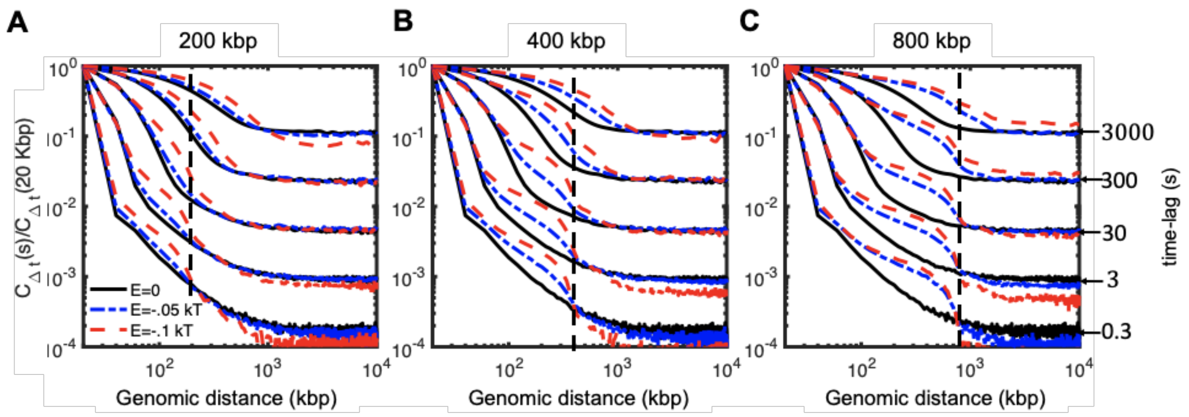


**Suppl. Fig. S12: Correlation of motions as a function of the genomic distance for uniform models with different strength of interactions, TAD lengths and time-lags.** All the correlations were computed with respect to the lab frame. The correlation length is increasing with the strength of interaction and with the time-lag. Increasing the compaction level of TADs leads to higher correlation of motions of intra-TAD loci.

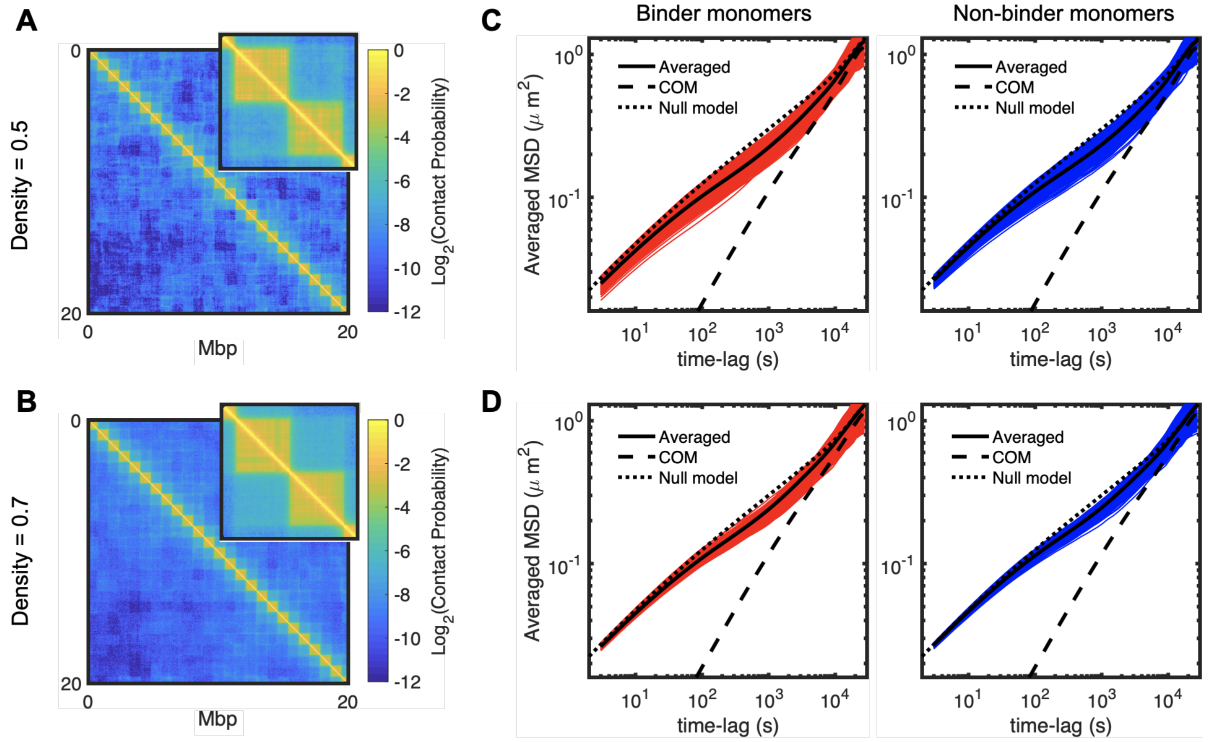


**Suppl. Fig S13: The effect of non-homogeneous models on dynamical heterogeneity and anomalous diffusion.** In contrast with the uniform model, we implemented a model where only a random subset of monomers inside the TAD can bind to each other. We varied the density $\rho$ of binder monomers and also the strength of interaction to have similar TAD compaction than in the uniform case with $E = -0.05kT$ (see Fig. 6B). **(A,B)** Predicted Hi-C maps (and corresponding 2Mb zooms) for $\rho = 0.5$ (A) and 0.7 (B). **(C,D)** Ensemble-averaged MSDs of all binder monomers (red) and non-binder, neutral monomers (blue) along with the averaged, center of mass and null model predictions. We found that such a model leads to heterogeneous dynamics (even higher than in the uniform models) and also to coherent motion and anomalous behavior. The effect is stronger for more compacted structures.

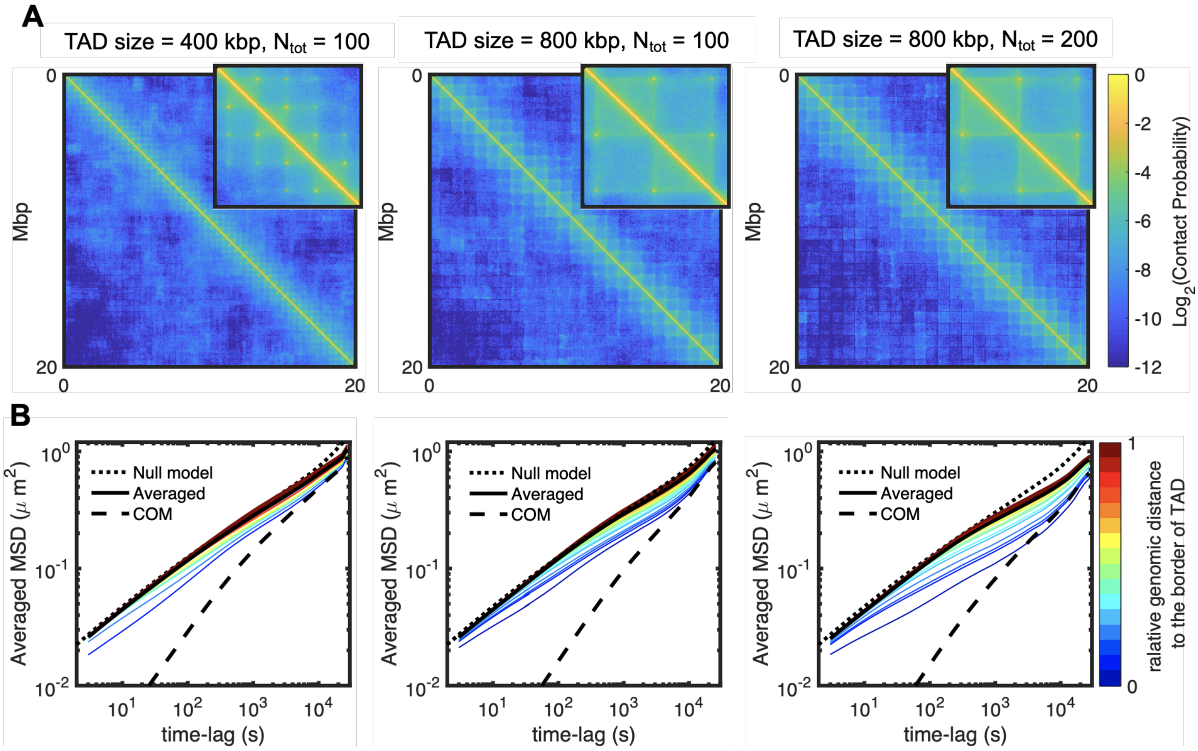


**Suppl. Fig S14: Effect of loop extrusion model on heterogeneity and anomalous diffusion.** We developed a toy, loop-extrusion model in which chromosomes are partitioned into similar size TADs. We explored the effect of TAD length and of the total number of extruding factors ($N_{tot}$) on chromosome mobility. **(A)** Predicted Hi-C matrices (and corresponding 2Mb zooms) for three different cases. **(B)** Ensemble-averaged MSDs of monomers inside TAD. The color code represents the relative location to the nearest TAD border (0 means at the border and 1 at the middle). The prediction for the null model and the MSD of the center of mass of the TAD are also shown. We found that the loop extrusion model leads to heterogeneous dynamics (even higher than in the uniform models) and also to coherent motion and anomalous behavior. The effect is stronger for more compacted structures.

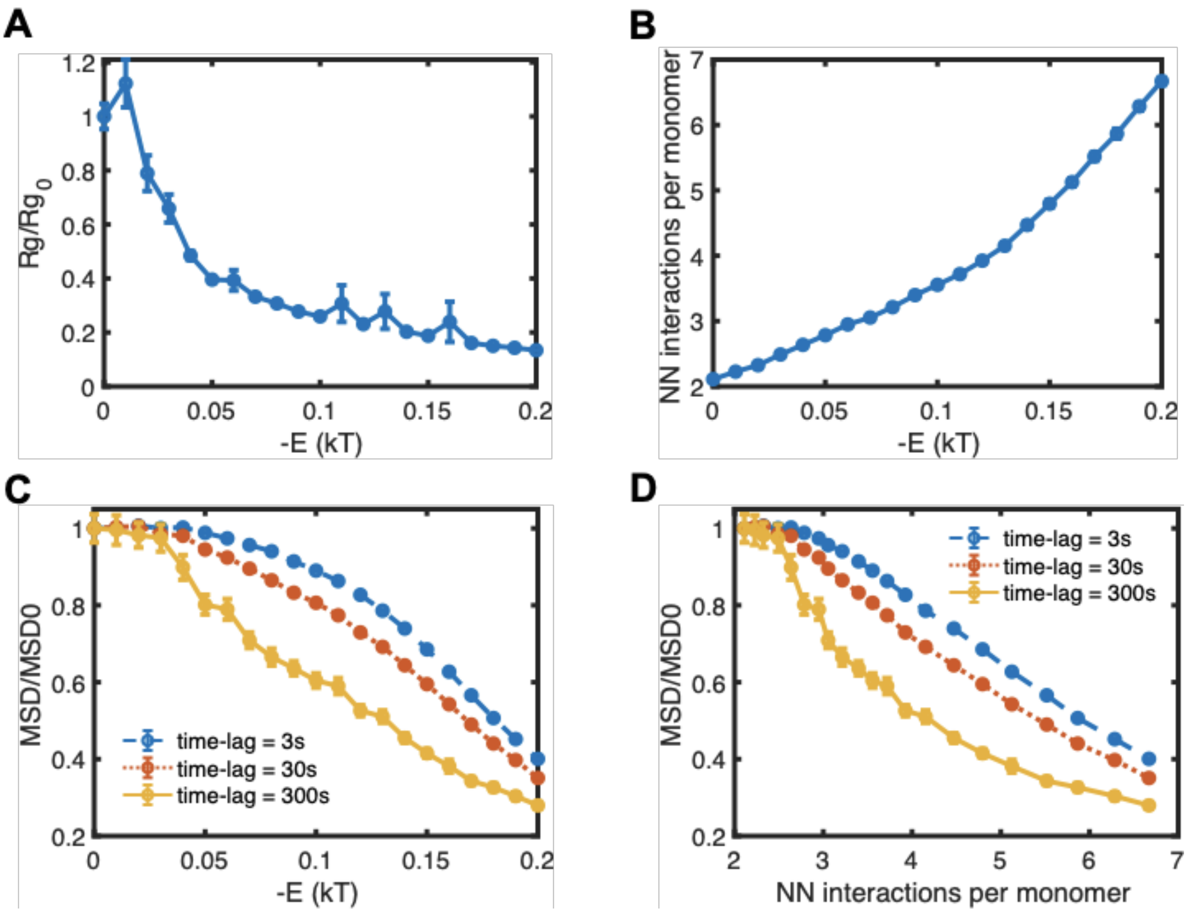


**Suppl. Fig S15: The effect of intra-TAD strength of interaction on compaction and dynamics.** For a toy, uniform model with 800kb-long TAD length (as in **Suppl. Fig. S3**), we varied the intra-TAD strength of interactions. (A) Compaction level (measured radius of gyration normalized by the corresponding radius of gyration in the null model) of TAD, as a function of strength of interaction. (B) Average number of nearest-neighbor (NN) contacts on the lattice between monomers of the same TAD. (C) Mean squared displacement for a given time-lag divided by the corresponding MSD in the null model, as a function of strength of interaction. (D) Normalized MSD as a function of the number of NN contacts. As the number of contact continuously increases in the system, TADs observe a structural, theta-collapse-like transition (A) and a smooth dynamical transition (D). These coupled transitions are characteristic of weak gelation in polymers with reversible crosslinks (de Gennes 1979).

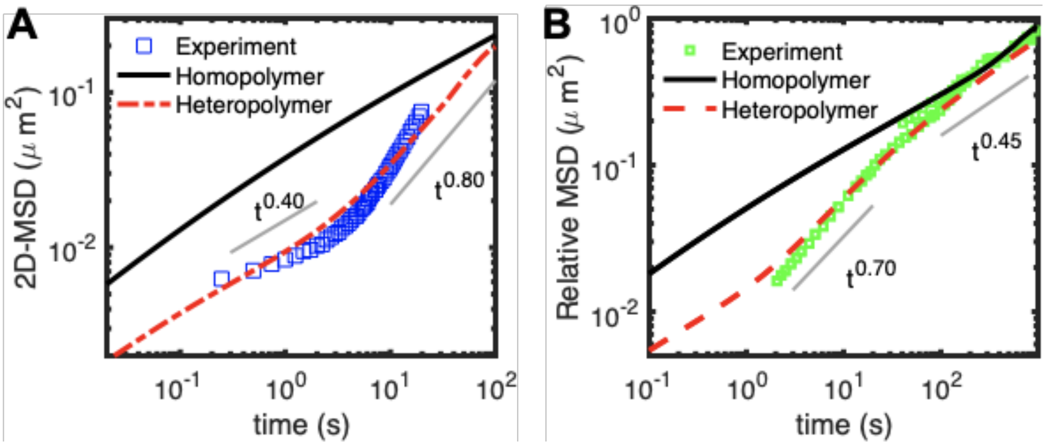


**Fig S16: Comparison with experimental data on the motion of single loci**. We compared the prediction of the heteropolymer model with two different experimental MSDs with anomalous behaviors that have been previously published. **(A)** Experimental data of the MSD of transcriptionally deactivated transgene in the human breast cancer cell line MCF-7 (Germier et al 2017). It exhibits a crossover between two different regimes with $\alpha < 0.5$ (sub-Rousean regime) at time-lags < 1s and $\alpha > 0.5$ (super-Rousean regime) at time-lags > 10s. This behaviour is qualitatively in good agreement with the heteropolymer by considering a 100 kbp region and fit the intra-TAD strength of interaction ($-0.3\ kT$) to reproduce the anomalous behavior. **(B)** Experimental data of the MSD observed at the Igh locus between the two alleles of the $D_HJ_H$ segment of pro-B cells (Khanna et al, 2019). The relative MSD is also anomalous with a super-Rousean regime at time-lags < 20s and a normal Rousean regime ($\alpha$ ~0.45) at larger time-lags. To compare with model predictions, we considered a 100 kbp region and fit the intra-TAD strength of interaction ($-0.2\ kT$) to reproduce the anomalous relative MSD between the two DJ alleles. To do that we assumed that the relative MSD is equal to the standard 2D MSD of one allele (as the motion of the two alleles, being on separated chromosomes, can be considered as uncorrelated). Time mapping for (A) and (B) was done as described in the **Methods** part of the main text.

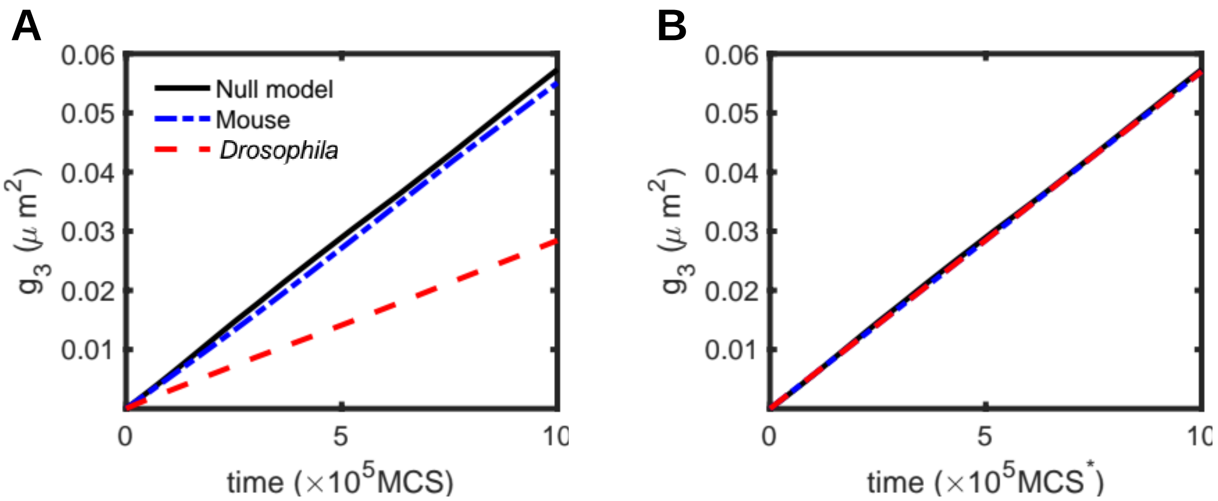


**Suppl. Fig. S17: Rescaling of MCS time steps. (A)** MSD of the center of mass ($g_3$) as a function of $MCS$ (A) and rescaled time $MCS^*$ **(B)** for homopolymer (black, full curves), mouse (blue, dash-dotted curves) and *Drosophila* (red, dashed curves).

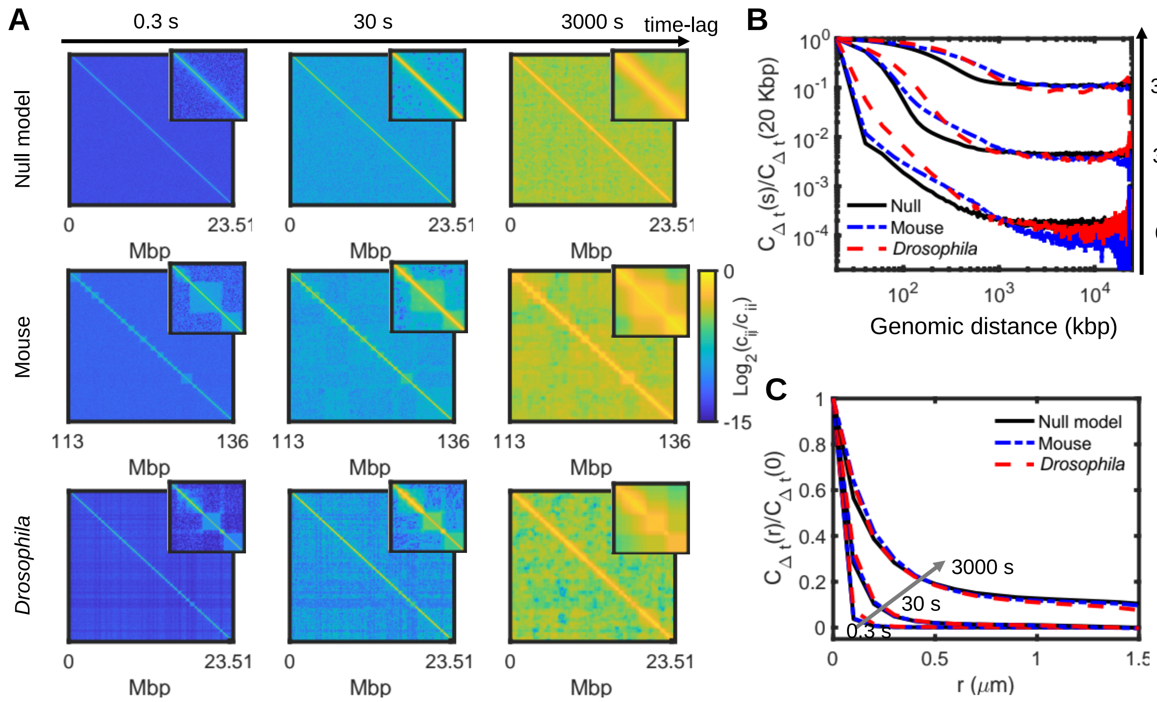


**Suppl. Fig S18: Spatiotemporal correlations of loci displacements with respect to lab frame. (A)** Normalized matrices of pair correlations without correcting the motion of the center of mass in the homopolymer (top panels), mouse (middle panels) and *Drosophila* (bottom panels) cases for different time-lags. Insets show 2 Mbp zoom into matrices. **(B,C)** Normalized correlations as a function of the genomic distance (B) and spatial distance (C) for the homopolymer (full), mouse (blue) and *Drosophila* (red) cases for different time-lags. Arrows indicate increasing time-lag.
